# Satellite glial cells promote regenerative growth in sensory neurons

**DOI:** 10.1101/2019.12.13.874669

**Authors:** Oshri Avraham, Pan-Yue Deng, Sara Jones, Rejji Kuruvilla, Clay F. Semenkovich, Vitaly A. Klyachko, Valeria Cavalli

## Abstract

Peripheral sensory neurons switch to a regenerative state after nerve injury to enable axon regeneration and functional recovery. Intrinsic mechanisms operating in sensory neurons are known to regulate nerve repair, but whether satellite glial cells (SGC), which completely envelop the neuronal soma, undergo injury-evoked transcriptional changes and contribute to nerve regeneration remains unexplored. This is largely due to the lack of molecular and genetic tools to study SGC. Using a single cell RNAseq approach to define the transcriptional profile of SGC in naïve and injured conditions, we reveal that these cells are distinct from Schwann cells and share similarities with astrocytes. We find that nerve injury elicits gene expression changes in SGC, which are related to fatty acid synthesis and peroxisome proliferator-activated receptor (PPARα) signaling. Conditional deletion of Fatty acid synthase (Fasn), the committed enzyme in *de novo* fatty acid synthesis, in SGC, impairs axon regeneration. The PPARα agonist fenofibrate rescues the impaired axon regeneration in mice lacking Fasn in SGC, indicating that PPARα functions downstream of fatty acid synthesis in SGC to promote axon regeneration. These results identify fatty acid synthesis in SGC as a fundamental novel mechanism mediating axon regeneration in adult peripheral nerves. These results also highlight that the sensory neuron and its surrounding glial coat form a functional unit that orchestrates nerve repair.

## INTRODUCTION

Unlike neurons in the central nervous system, peripheral sensory neurons with cell soma in dorsal root ganglia (DRG) switch to a regenerative state after nerve injury to enable axon regeneration and functional recovery. Decades of research have focused on the signaling pathways elicited by injury in sensory neurons ^1, 2^ and in Schwann cells that insulate axons ^3, 4^ as central mechanisms regulating nerve repair. However, virtually nothing is known about the contribution of the glial cells that envelop the neuronal soma, known as satellite glial cells (SGC), to the nerve repair process.

In adult animals, multiple SGC form an envelope that completely enwraps each sensory neuron soma ^5, 6, 7^. The number of SGC surrounding sensory neurons soma increases with increasing soma size in mammals ^8, 9^. Each sensory neuron soma and its surrounding SGC are separated from adjacent neurons by connective tissue ^5, 6, 7^. The neuron and its surrounding SGC thus form a distinct morphological unit ^5, 6, 9^. Structural neuron-glia units similar to these do not exist in the central nervous system ^5, 6, 7^. SGC have been identified mostly based on their morphology and location. Several SGC markers have been characterized, including the inwardly rectifying potassium channel (Kir4.1) ^10^, cadherin 19 (CDH19) ^11^, the calcium activated potassium channel (SK3) ^12, 13^ and glutamine synthetase (GS) ^7, 14^. SGC also share several properties with astrocytes, including expression of glial fibrillary acidic protein (GFAP) ^7, 15, 16^ and functional coupling by gap junctions ^7, 16, 17^.

SGC have been studied in the context of pain responses and are known to modulate pain thresholds ^18, 19, 20, 21, 22^. It is known that SGC are altered structurally and functionally under pathological conditions, such as inflammation, chemotherapy-induced neuropathic pain and nerve injuries ^7, 22, 23, 24^. Communication between neuron soma and SGC via glutamatergic transmission could impact neuronal excitability and thus nociceptive threshold after injury ^25^. Nerve lesions induce an increase in GFAP ^16, 22, 26, 27, 28, 29^, p75 and pERK expression ^30, 31^ in SGC. An increase in SGC ongoing cell division following nerve injury has also been suggested ^16, 32^.

These studies suggest that SGC can sense and respond to a distant nerve injury and actively participate in the processing of sensory signals ^33^. Whether SGC play a role in regenerative responses has not yet been established. In this study, using single cell RNA-seq, we reveal that nerve injury alters the gene expression profile in SGC, which is mostly related to lipid metabolism, including fatty acid synthesis and PPARα signaling.

PPARs are ligand-activated nuclear receptors with the unique ability to bind lipid signaling molecules and transduce the appropriate signals derived from the metabolic environment to control gene expression ^34^. After binding the lipid ligand, PPARs form a heterodimer with the nuclear receptor RXR, followed by binding to specific DNA-response elements in target genes (PPAREs). Three different PPAR subtypes are known; PPARα, PPARβ/δ and PPARγ ^35^. The committed enzyme in *de novo* fatty acid synthesis Fatty acid synthase (Fasn),^36^ generates endogenous phospholipid ligands for PPARα ^37^ and PPARγ ^38^. PPARγ activity has been associated with neuroprotection in different neurological disorders ^39^. In rat sensory axons, PPARγ contribute to the pro-regenerative response after nerve injury ^40^. In the CNS, astrocytes produce lipids far more efficiently than neurons and Fasn was found in astrocytes but not in neurons ^41^. Lipids secreted in ApoE-containing lipoproteins by glial cells appear to support growth in cultured hippocampal neurons and regulate expression or pro-regenerative genes such as Gap43 ^42^. In the PNS, synthesis of phospholipids and cholesterol in sensory neurons is required for axonal growth ^43, 44, 45, 46^. But lipids can also be exogenously supplied by lipoproteins secreted from glial cells to stimulate neurite growth ^42, 45, 47, 48^, possibly via ApoE, whose expression is increased in glial cells after nerve injury ^49, 50^. Whether SGC express PPAR or Fasn and contribute to support sensory axon growth after nerve injury has not been determined.

Here, we demonstrate that conditional deletion of Fasn specifically in SGC impairs axon regeneration in peripheral nerves. Treatment with fenofibrate, an FDA-approved PPARα agonist, rescues the impaired axon regeneration observed in mice lacking Fasn in SGC, suggesting that PPARα functions downstream of fatty acid synthesis in SGC to promote axon regeneration. These results unravel lipid synthesis in SGC as a fundamental novel mechanism mediating axon regeneration in adult peripheral nerves. These results also highlight that the neuron and its surrounding glial coat form a functional unit that orchestrates nerve repair.

## RESULTS

### Single-cell transcriptional profiling identified multiple cell types in naïve and injured DRG

To define the biology of SGC and to understand the role of SGC in nerve injury responses, we performed single cell RNA-seq (scRNA-seq) of mouse L4,L5 DRG in naïve and injured conditions (3 days post sciatic nerve crush injury), using the Chromium Single Cell Gene Expression Solution (10x Genomics) (Fig. 1a). An unbiased (Graph-based) clustering, using Partek flow analysis package, identified 13 distinct cell clusters in the control and injury samples (Fig. 1b). The number of sequenced cells was 6,541 from 2 biological replicates, with an average of 45,000 reads per cell, 1,500 genes per cell and a total of 17,879 genes detected (see filtering criteria in the methods). To identify cluster-specific genes, we calculated the expression difference of each gene between that cluster and the average in the rest of the clusters (ANOVA fold change threshold >1.5), illustrated by a heatmap of the top 5 differentially expressed genes per cluster relative to all other clusters (Fig. 1c and Table 1). Examination of the cluster-specific marker genes revealed major cellular subtypes including neurons (Isl1), SGC (KCNJ10/Kir4), Schwann cells (Periaxin, MPZ), endothelial cells (Pecam1/CD31), macrophages (Alf1/Iba-1, CD68), mesenchymal (CD34) connective tissue (Col1A1), T-cells (CD3G) and Vasculature associated smooth muscle cells (DES) (Fig. 1b and S1a). We then compared the cell clustering between naïve and injury conditions. Unique cell clusters were not altered by nerve injury (Fig. 1d). We found that the number of SGC and macrophage were increased after injury by 8% and 7%, respectively (Fig. 1e). Although prior studies suggested that SGC proliferate after injury ^7, 13, 16, 32^, a recent study demonstrated that cell proliferation occurred in macrophages but not in SGC seven days after injury ^51^. We thus examined expression of the cell cycle markers MKI67 and CDK1 in injury conditions and found that these cell cycle markers were mainly expressed in macrophages and blood cells/monocytes but not in SGC (Fig. S1b). Our results thus suggest that 3 days post injury, there is little SGC proliferation and the increase in SGC cell number we observed could be a result of tissue dissociation. We obtained a similar number of neurons in both uninjured and injured conditions, which represented about 1% of high quality sequenced cells (Fig. 1e). The recovered neurons included nociceptors (TrkA), mechanosensors (TrkB) and proprioceptors (TrkC) (Fig. S1c). We anticipated obtaining a larger representation of neurons in our dataset. The dissociation procedure, which requires multiple steps of enzymatic and mechanical disruption to separate the SGC from neurons might affect neuronal survival. Our protocol is thus achieving recovery of SGC, but might not be suitable for the recovery of neurons from DRG.

**Figure 1:**
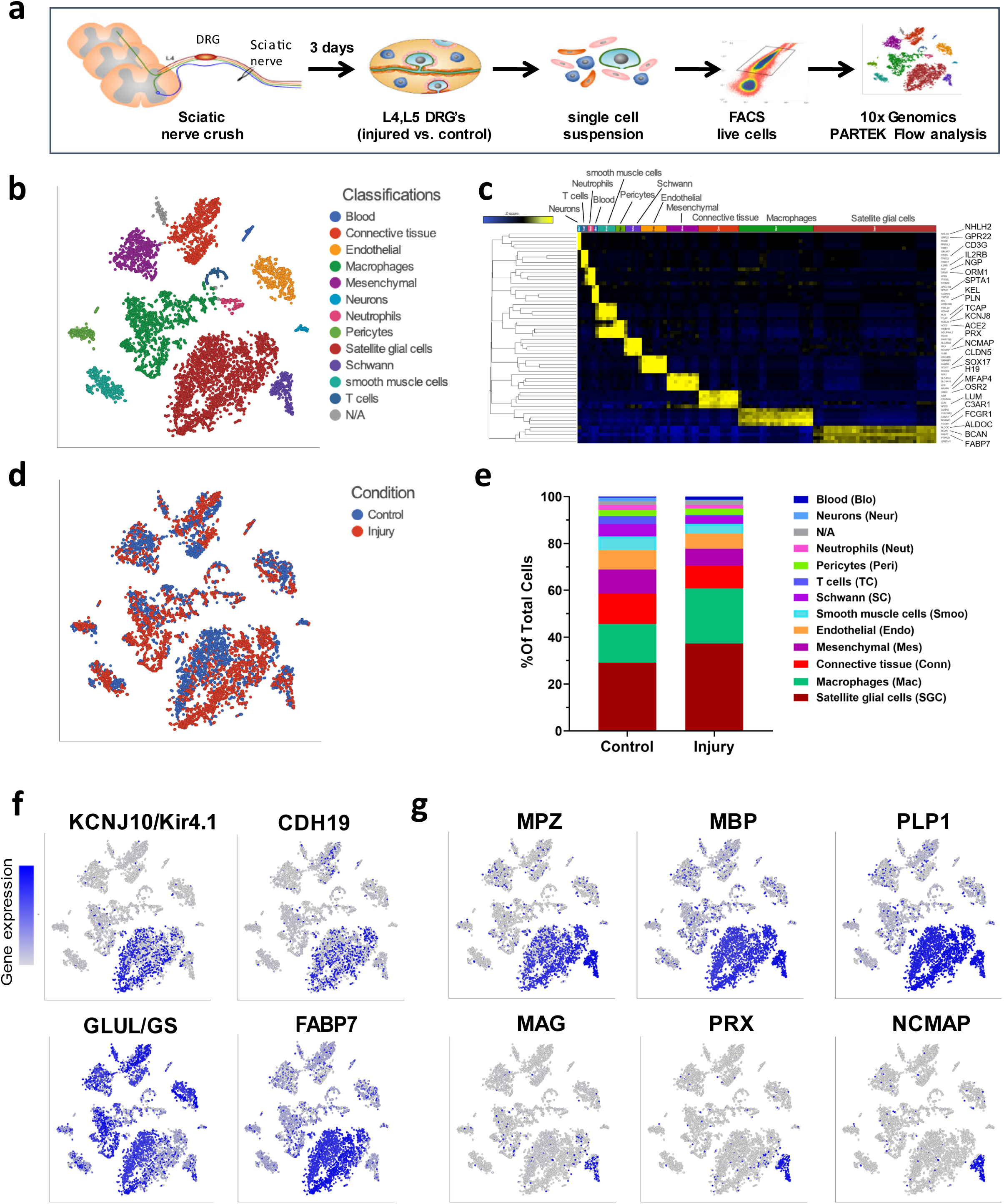
Characterization of cell populations in the DRG. (a) Schematic of the experimental design for scRNAseq (b) t-SNE plot of 6,541 cells from L4,L5 mouse dissociated DRG, with 13 distinct cell clusters (Unbiased, Graph based clustering). Classifications were assigned based on known marker genes. (c) Heatmap of the top five differentially expressed genes per cluster relative to all other clusters. (d) t-SNE plot of cells from naïve (blue) and injury (red) conditions. (e) Fraction of each cell type within control (2,915 cells) and injury (3,626 cells) conditions. Source data are provided as a Source Data file (f) t-SNE overlaid for expression of marker genes for SGC (g) t-SNE overlaid for expression of marker genes shared between Schwann cells and SGC (top panel) and marker genes for Schwann cell only (bottom panel).

### SGC are molecularly distinct from Schwann cells in DRG

The scRNAseq results indicate that SGC represent the largest glial subtype in the DRG (Fig. 1e). Overlaid cells in t-SNE plots with marker genes for SGC and Schwann cells show the relative levels of expression as a color gradient (Fig. 1f and 1g). The SGC cluster identity was confirmed by the expression of known SGC markers such as Kir4.1 ^10^ and CDH19 ^11^ (Fig. 1f and S1d). Whereas glutamine synthetase (GLUL/GS) has been used as a SGC specific marker at the protein level ^51, 52^, our analysis showed a nonspecific expression in almost all cells in the DRG (Fig. 1f and S1d). Another marker used for SGC in rat is SK3 ^52^, but SK3 was not detected in mouse SGC nor any other cells in the DRG. The most highly expressed gene in SGC, Fatty Acid Binding Protein 7 (Fabp7, also known as BLBP and BFABP) was enriched in SGC (Fig. 1f, S1d and Table 1). Whereas SGC and Schwann cells share the expression of known Schwann cell markers, such as MPZ, MBP and PLP1 (Fig. 1g and S1e) the Schwann cell marker genes Myelin Associated Glycoprotein (MAG), Periaxin (PRX) and Non-Compact Myelin Associated Protein (NCMAP) genes were not expressed in SGC (Fig. 1f and S1e). We next compared the top differentially expressed genes in SGC (605 genes) and Schwann cells (572 genes) (ANOVA threshold >4 fold change p-value<0.05 compared to all other populations in the DRG), which revealed that SGC share only 2% of those gene transcripts with Schwann cells in the DRG (Fig. S1f and Table 2). A recent study highlighted the molecular differences between myelinating and non-myelinating Schwann cells in the sciatic nerve ^53^. Comparison of the SGC molecular profiles revealed higher similarity between SGC and myelinating Schwann over non-myelinating Schwann cells (Table 2). SGC also share several properties with astrocytes, including expression of Kir4.1 and glial fibrillary acidic protein (GFAP) ^7, 10, 15^. We thus examined the transcriptional similarity between astrocytes and SGC by comparing the top differentially expressed genes in each cell type, using our scRNAseq data for SGC (605 genes, >4 fold change p-value<0.05 compared to all other populations in the DRG) and a previously published transcriptional analysis of astrocytes (500 genes, >6 fold change compared to other populations in the cerebral cortex) ^54^. We found that SGC share about 10% of those gene transcripts with astrocytes, among them Fabp7, Gfap, Ppara and Aldoc (Fig. S1g, Table 2). Our single cell RNAseq analysis using freshly dissociated tissue thus unravels the unique transcriptional profile of SGC in DRG and reveals that Schwann cells and SGC are transcriptionally distinct in the DRG.

### Fatty acid binding protein 7 is a specific marker for adult mouse SGC

The scRNAseq data revealed that one of the top differentially expressed genes in the SGC cluster is Fabp7 (Fig. 1f and Table 1). Fabp7 is a nervous system specific member of the hydrophobic ligand binding protein family involved in uptake and transport of fatty acid ^55^. Fabp7 is involved in signal transduction and gene transcription in the developing mammalian CNS ^56^, but its precise function remains quite elusive. In the peripheral nervous system, Fabp7 is expressed during embryonic development and distinguishes glia from neural crest cells and neurons during the early stages of development ^57, 58, 59 60^. These observations prompted us to further test if Fabp7 can be used as a novel marker of SGC in adult mouse DRG. Consistent with our single cell data, we found that Fabp7 labels SGC surrounding sensory neuron soma (Fig. 2a). The specificity of the Fabp7 antibody was verified using DRG and sciatic nerve from the Fabp7KO mouse ^61^ (Fig. S2a-e), in which Fabp7 signal surrounding neurons is lost, but SGC are present and stained with glutamine synthetase (GS) ^7, 14^. Importantly, Fabp7 does not label Schwann cells surrounding axons in the DRG (Fig. 2a, asterisks), or in the sciatic nerve (Fig. S2d,e) consistent with previous reports ^62, 63^ and our single cell analysis (Fig. 1f). These results suggest that Fabp7 protein expression is highly enriched in SGC in the DRG.

**Figure 2:**
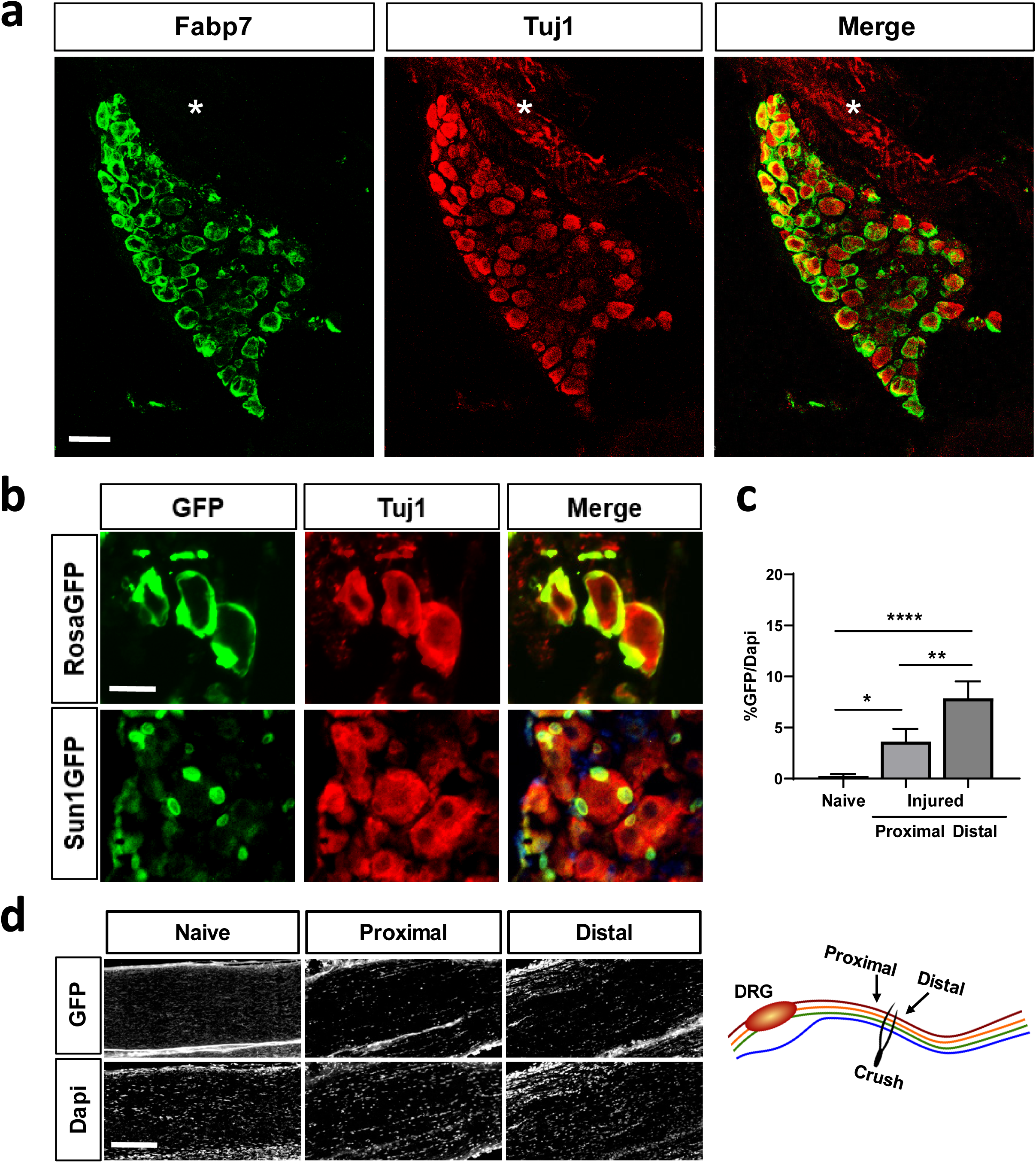
Fabp7 is a specific marker for SGC. (a) Representative images of immunofluorescence staining of mouse DRG sections with Fabp7 (green) which labels SGC surrounding neurons marked with Tuj1 (red). No Fabp7 expression in the axon rich area (asterisks) is observed. n=3, Scale bar: 100 µM. (b) BlbpCre-ER mice crossed with RosaGFP or Sun1GFP show expression of GFP in the SGC surrounding the neurons marked with Tuj1 (red). RosaGFP n=3, Sun1GFP n=4 Scale bar: 50 µM. (c) Quantification of Sciatic nerves from BlbpCre-ER mice crossed with Sun1GFP. GFP positive nuclei normalized to the total number of dapi positive nuclei. One-way analysis of variance (ANOVA) Sidak’s multiple comparisons test. *p<0.05 **p<0.005; **** p<0.0001. Source data are provided as a Source Data file (d) Represented images from naive and injured nerves, ∼0.5 mm proximal and distal to the injury site as shown in the scheme. n=4 Scale bar: 50 µm.

We next tested if the BLBPcre-ER mouse line ^64^ can be used to label and manipulate SGC specifically. We crossed BLBPCre-ER to the Rosa26-fs-TRAP and observed that following a 10 days tamoxifen treatment, BLBPcre-ER mice drove expression of the GFP reporter in SGC (Fig. 2b, upper panel). To further ensure that the BLBPcre-ER is specifically expressed in SGC in the DRG and does not drive expression in Schwann cells in the nerve, we crossed BLBPcre-ER to the Sun1-sfGFP-myc (INTACT mice: R26-CAG-LSL-Sun1-sfGFP-myc) ^65^, which allows GFP expression in the nuclear membrane. We found GFP positive nuclei around the neurons in DRG sections (Fig. 2b, lower panel) and no nuclear GFP expression axons in naïve sciatic nerves (Fig. 2c,d). Since a role for Fabp7 in regulating Schwann cell-axon interactions has been proposed ^62^, with Fabp7 expressed 2 to 3 weeks after nerve injury when Schwann cell process formation is exuberant, we also examined nuclear GFP expression in injured nerves. We found that 3 days post injury, 4% and 7% of the nuclei expressed the GFP reporter approximately 0.5mm proximal and distal to the injury site, respectively (Fig. 2c,d). This is consistent with a prior report showing that BLBP is not present 4 to 7 days post injury, but is expressed in some cells starting 14 days post injury ^62^. Since typically 50% of nuclei express c-jun, a marker for Schwann cells response to injury, two days after injury in the nerve ^66^, the GFP positive nuclei we observed, may represent a very small subset of Schwann cells responding to injury. To further confirm the specificity of the BLBPCreER reporter following nerve injury, we re-analyzed a single cell data set from injured nerve (9 days post injury) ^67^. We found that less than 5% of cells in the Schwann cell cluster expressed Fabp7 (Fig. S2f). A comparative analysis of the transcriptome of injured nerve (3 days post injury) also demonstrated very low counts of Fabp7 in proximal and distal nerve segments ^68^. Because Fabp7 is also expressed at a low level in a subset of other cell types including macrophages, endothelial cells and mesenchymal cells (Fig. 1f and S1e), we cannot rule out the possibility that BLBPCreER may also label a subset of other cells in the DRG. Together, these experiments reveal that Fabp7 represents a novel marker of SGC and that the BLBPcre-ER mouse line can be used to label and manipulate SGC, with only minimal impact on Schwann cells at early time points after nerve injury.

### SGC upregulate lipid metabolism in response to nerve injury

To define the transcriptional response of SGC to nerve injury, cells in the SGC cluster were pooled, control and injury conditions were compared to identify differentially expressed genes. 1,255 genes were differentially upregulated in SGC after injury (FDR>0.05, Log2Fold change>2) (Table 3) and analyzed for enriched biological pathways using KEGG 2016 (Kyoto Encyclopedia of Genes and Genomes). This analysis revealed enrichment in lipid metabolic pathways, including fatty acid biosynthesis (Fig. 3a). Our findings also confirmed the upregulation of previously reported injury induced genes in SGC such as Connexin43/GJA1 and GFAP ^16, 23^ (Fig. S3g, Table 3). 300 genes were differentially downregulated in SGC, with enrichment for cell cycle and p53 signaling pathways (Table 3 and 4). Pathway analysis of upregulated genes in other major cell types in the DRG confirmed fatty acid metabolism as a unique pathway enriched in SGC in response to nerve injury (Fig. S3a-f). This analysis also indicated that the cell cycle term was enriched in macrophages, and downregulated in SGC (Fig. 3a and S3a, Table 4), supporting the recent finding that macrophages but not SGC undergo cell cycle after nerve injury^51^.

**Figure 3:**
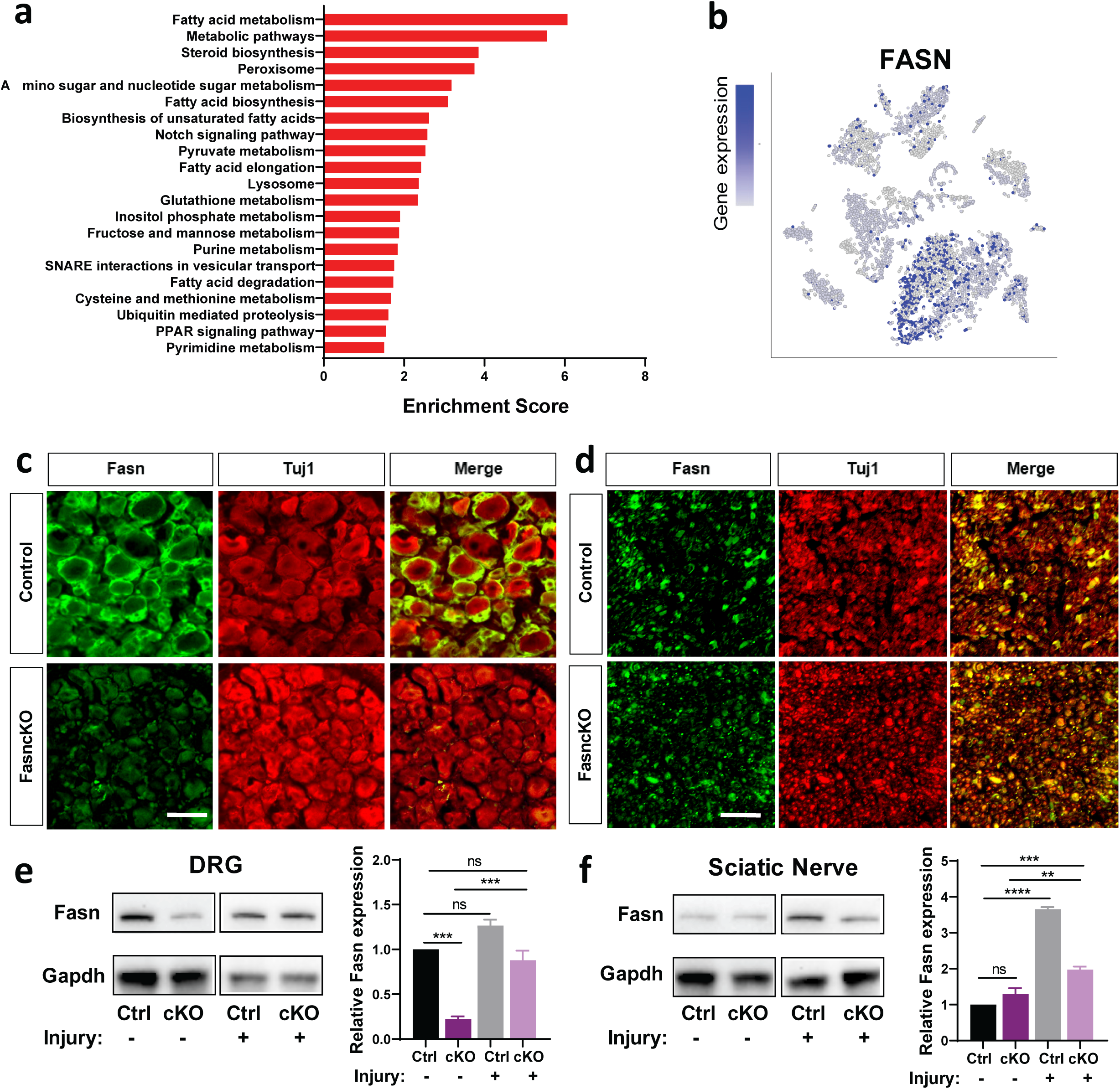
SGC upregulate genes involved in lipid metabolism in response to nerve injury. (a) Pathway analysis (KEGG 2016) of differentially upregulated genes in the SGC cluster. (FDR<.05, Log2Fold change>2) (b) t-SNE overlay for expression of Fasn gene. (c) DRG sections from control and FasncKO mice, immunostained for Fasn (green) and the neuronal marker Tuj1 (red) n=3. C, Scale bar: 50 µm (d) Nerve sections from control and FasncKO mice, immunostained for Fasn (green) and the neuronal marker Tuj1 (red) n=3. C, Scale bar: 50 µm. (e) Western blot analysis and quantification of Fasn protein expression in DRG from control and FasncKO mice with and without injury. Quantification of Fasn expression normalized to Gapdh expression. n=3. One way ANOVA. Dunnett’s multiple comparisons test. *** p<0.001 ns-non significant. Source data are provided as a Source Data file (f) Western blot analysis and quantification of Fasn protein expression in Sciatic nerve from control and FasncKO mice with and without injury. Quantification of Fasn expression normalized to Gapdh expression. n=3. One way ANOVA. Dunnett’s multiple comparisons test. **** p<0.0001, *** p<0.001, ** p<0.01 ns-non significant. Source data are provided as a Source Data file

One of the genes enriched in SGC is fatty acid synthase (Fasn), which controls the committed step in endogenous fatty acid synthesis ^36^ (Fig. 3b and Table 2). Although Fasn levels were not differentially regulated after injury in SGC, we decided to explore the role of Fasn further, because Fasn has the potential to impact major signaling and cellular pathways ^69^. Fasn product is converted to other complex lipids that are critical for membrane structure, protein modification and localization ^69^ and are also utilized for phospholipid synthesis. Immunostaining of DRG sections confirmed Fasn expression in SGC surrounding sensory neuron soma (Fig. 3c, upper panel). To investigate the role of Fasn in SGC, we generated an SGC specific *Fasn* KO mouse (FasncKO) by crossing BLBPcre-ER ^64^ mice to mice carrying floxed *Fasn* alleles ^70^. Eight week old BLBPcre-ER ;Fasn^f/f^ (FasncKO) and BLBPcre-ER^-^ ;Fasn^f/+^ mice (control) were fed with tamoxifen for 10 days to conditionally delete Fasn in SGC. This design allows us to ensure that both control and FasncKO groups are treated with tamoxifen and thus differences between groups do not result from the tamoxifen treatment itself. Expression of Fasn in SGC visualized by immunofluorescence was significantly reduced in the FasncKO DRG compared to control littermates (Fig. 3c, bottom panel). Western blot analysis of Fasn in DRG also confirmed a significant reduction of Fasn expression in the FasncKO compared to control littermates (Fig. 3e). Fasn has been shown to be expressed in Schwann cells ^71^, and immunostaining for Fasn in the sciatic nerve support these results (Fig. 3d). However, Fasn expression was not affected in the nerve of FasncKO mice (Fig. 3d and 3f) supporting the specificity of the BLBPCre-ER line for SGC. We also examined Fasn protein expression in DRG and sciatic nerve by western blot in naive and injured conditions. Nerve injury increased the protein level of Fasn in the DRG of FasncKO but not control mice (Fig. 3e). Nerve injury also increased Fasn levels in the nerve of both control and FasncKO (Fig. 3f) which could reflect an increase in Fasn levels in Schwann cells after nerve injury ^68, 71^. Together, these results indicate that Fasn can be efficiently deleted from SGC in the DRG, with minimal impact in Schwann cells.

### Deletion of Fasn in SGC does not alter neuronal morphology or their functional properties

To determine the impact of Fasn deletion on SGC and neuron morphology, we performed transmission electron microscopy (TEM) as previously reported ^6, 9^. SGC surrounding the neuron soma were pseudo colored (Fig. 4a), We observed an increase in the SGC nuclear area and a more circular, less elliptic nuclear morphology in the FasncKO animals compared to controls (Fig. 4a-c), whereas there was no change in the neuronal nucleus circularity (Fig. 4d). It has been reported that under certain conditions, such as alterations in lipid composition, the overall nuclear structure can be modified ^72^. A study in yeast demonstrated that deletion of certain genes affecting lipid biosynthesis leads to nuclear expansion ^72^, suggesting that Fasn deletion in SGC may affect nuclear morphology. However, in both control and FasncKO, the SGC sheath is smooth and separated from the sheaths enclosing the adjacent nerve cell bodies by a connective tissue space (Fig. 4a), as described extensively previously ^6, 9^. The SGC coat is also in direct contact with the neuronal membrane (Fig. 4a), as previously described ^6, 9^. To ensure that Fasn deletion in SGC does not impact nerve morphology, we next evaluated both Remak bundle structure and myelin sheath thickness in the nerve of the FasncKO mice compared to control animals (Fig. 4e-i). Neither the number of axons per Remak bundle nor the axon diameter and myelin thickness, measured as the ratio between the inner and outer diameter of the myelin sheath (g-ratio), was altered in the FasncKO. The overall nerve structure examined by TUJ1 staining was also not altered in the FasncKO nerves compared to controls in naïve conditions or following an injury (Fig. S4a).To ensure that the reduction of Fasn in SGC does not cause neuronal cell death, we immunostained sections of DRG from FasncKO and control mice for the apoptotic marker cleaved caspase 3 before and after injury (Fig. S4b). Quantification of Caspase 3 intensity revealed no change in apoptotic cell death in naïve conditions and three days post injury between FasncKO and control (Fig. 4j and S4b). These experiments do not exclude the possibility that other forms of cell death such as necrosis may occur.

**Figure 4:**
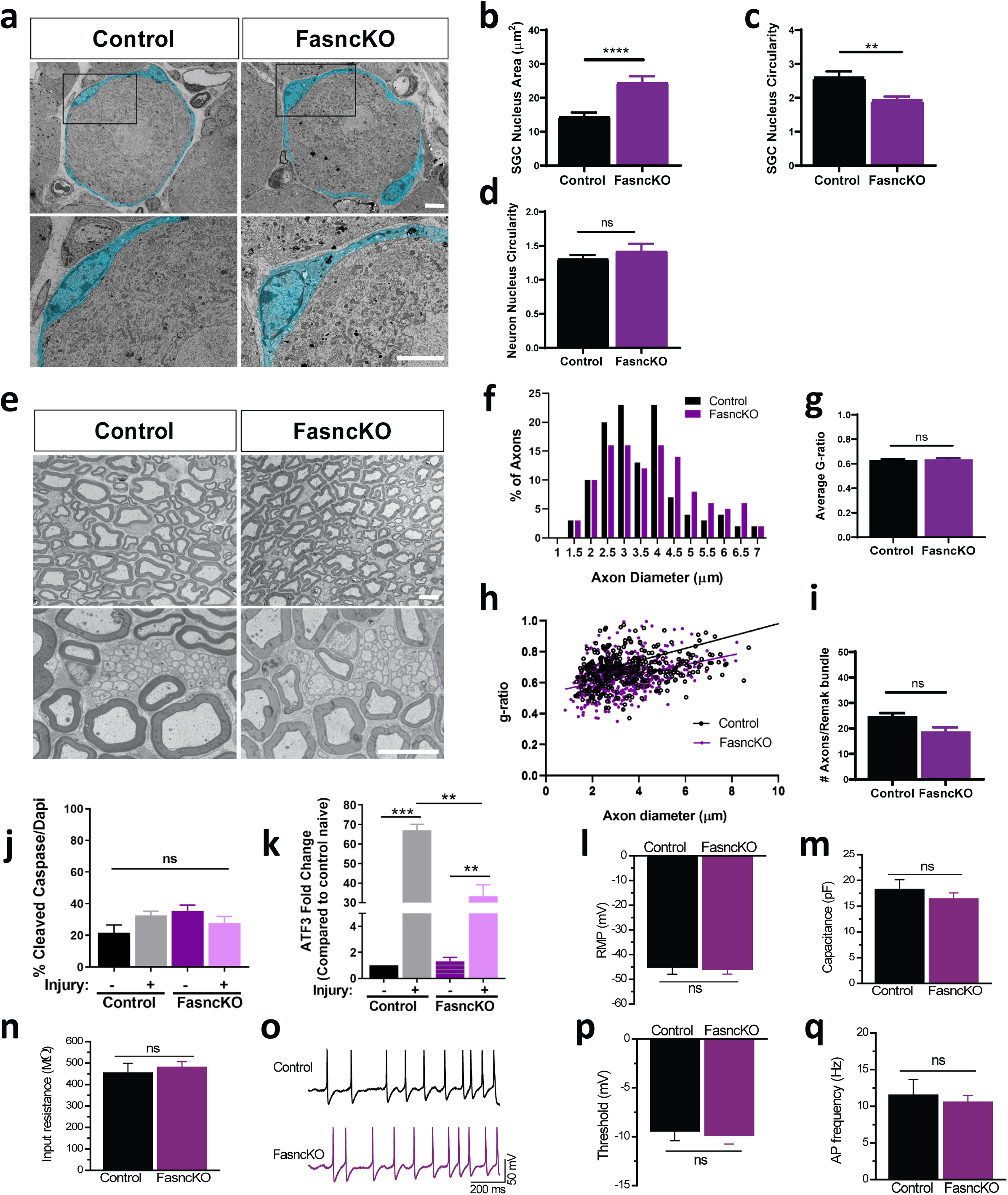
Fatty Acid synthase deletion in SGC does not alter neuronal morphology or functional properties. (a) Representative TEM images of a DRG cell body and its enveloping SGC sheath, pseudo-colored in turquoise, from control and FasncKO mice. High magnification of boxed area from top panels are showed in bottom panels. n=3 Scale bar: 5 µm (b) Average SGC nuclear area. t-test **** p<0.0001. Source data are provided as a Source Data file (c) SGC nuclear circularity (ratio between major axis(X) and minor axis(Y). 1=circular, 1<elliptic). t-test **p<0.005. Source data are provided as a Source Data file (d) Neuron nuclear circularity (ratio between major axis(X) and minor axis(Y). 1=circular, 1<elliptic). t-test ns-non significant. Source data are provided as a Source Data file (e) Representative TEM images of sciatic nerve cross sections from control and FasncKO mice. Scale bar: 5 µm. (f) Quantification of axon diameter distribution in sciatic nerves of FasncKO and control mice (average axon diameter Control=3.574 µm FasncKO=3.353 µm n=130 axons). Source data are provided as a Source Data file (g) Average g-ratio in sciatic nerves of FasncKO and control mice. t-test non-significant. Source data are provided as a Source Data file (h) Linear correlation of g-ratio versus axon diameter in sciatic nerves of FasncKO mice compared with controls. n=3 Unpaired t-test non-significant. Source data are provided as a Source Data file (i) Quantification of the number of axons per Remak bundle in FasncKO and control nerves. n=3 Unpaired t-test ns-non significant. Source data are provided as a Source Data file (j) Quantification of cleaved caspase 3 in DRG sections from control and FasncKO mice in naïve and 3 days after sciatic nerve injury. Ratio of Cleaved caspase3 positive vs. dapi was measured. n=4 One way ANOVA ns-non significant. Source data are provided as a Source Data file (k) qPCR analysis of ATF3 expression in DRG from control and FasncKO mice in naïve and 3 days after sciatic nerve injury n=3 One way ANOVA. Sidak’s multiple comparisons test **p<0.005 ***p<0.0005. Source data are provided as a Source Data file (l) Whole-cell recordings were performed in dissociated co-cultures of DRG neurons surrounded by glia from FasncKO and control mice. Quantification of resting membrane potential (RMP) was measured in control -45.3 ± 2.7 mV, n = 16; FasncKO -46.1 ± 1.8 mV, n = 30; p = 0.79; t-test,. Source data are provided as a Source Data file (m) Membrane capacitance (control 18.3 ± 1.8 pF, n = 16; FasncKO 16.5 ± 1.1 pF, n = 30; p = 0.36; t-test,. Source data are provided as a Source Data file (n) Input resistance (control 456 ± 43 MΩ, n = 16; FasncKO 482 ± 24 MΩ, n = 30; p = 0.56; t-test,. Source data are provided as a Source Data file (o) Spiking properties of DRG neurons from controls and FasncKO. (p) Potential threshold (control -9.46 ± 0.94 mV, n = 13; FasncKO -9.91 ± 0.84 mV, n = 28; p = 0.77 t-test, ns-non-significant. Source data are provided as a Source Data file (q) Neuronal firing rate (control 11.57 ± 2.09 Hz, n = 14; FasncKO 10.62 ± 0.86 Hz, n =29; p = 0.62 was measured. t-test, ns-non significant. Source data are provided as a Source Data file

We next examined the expression of ATF3, an established intrinsic neuronal injury marker ^1, 73^. ATF3 levels in control and FasncKO DRG were very low in the absence of injury (Fig. 4k), indicating that in naïve condition, Fasn deletion in SGC did not cause a stress response in DRG neurons. Nerve injury increased the mRNA levels of ATF3 in control and FasncKO DRG, but this response was less robust in the FasncKO (Fig. 4k). Together, these results indicate that loss of Fasn in SGC does not lead to neuronal morphological deficits in the DRG and does not elicit a stress response in neurons.

The bidirectional communication between neurons and SGC participates in neuronal excitability and the processing of pain signals ^33^. The excitability of sensory neurons is controlled in part by the surrounding SGC ^18, 74, 75, 76, 77^. To examine whether Fasn deletion in SGC affected functional properties of DRG neurons, we compared intrinsic excitability and firing properties of DRG neurons from FasncKO and control mice. Whole-cell recordings were performed in short-term cultures of DRG neurons and glia. Medium diameter neurons that were associated with at least one glial cell were targeted for recordings (Fig. S4c). A subset of recorded cells was filled with biocytin via the patch pipette for *post hoc* verification of neuronal identity. The vast majority of filled cells were identified as IB4-positive nociceptors (Fig. S4d). We observed that all major features of intrinsic neuronal excitability were unaffected in DRG neurons from FasncKO mice compared to controls, including the resting membrane potential (Fig. 4l), membrane capacitance (Fig. 4n), and input resistance (Fig. 4n). We further examined the spiking properties of DRG neurons and found no detectable changes in action potential threshold (Fig. 4p), which represents the principle determinant of neuronal excitability, or the neuronal firing frequency (Fig. 4o,c) in FasncKO mice compared to controls. Therefore, Fasn deletion in SGC does not affect functional properties of DRG neurons in the naïve, dissociated conditions.

### Deletion of Fasn in SGC impairs axon regeneration

We observed that ATF3 expression after injury was reduced in FasncKO compared to control (Fig. 4k), suggesting that absence of Fasn in SGC may impact the neuronal response to injury. To test the consequence of Fasn deletion in SGC on axon regeneration, we used our established *in vivo a*nd ex-vivo regeneration assays ^78, 79, 80^. Two weeks after tamoxifen treatment was completed, we performed a sciatic nerve crush injury in FasncKO and control mice and measured the extent of axon regeneration past the injury site three days later by labeling nerve sections with SCG10, a marker for regenerating axons ^81^. The crush site was determined according to highest SCG10 intensity along the nerve. We used two measurements to quantify axon regeneration. First, we measured the length of the 10 longest axons, which reflect the extent of axon elongation, regardless of the number of axon that regenerate. Second, we measured a regeneration index by normalizing the average SCG10 intensity at distances away from the crush site to the SCG10 intensity at the crush site. This measure takes into account both the length and the number of regenerating axons past the crush site. Loss of Fasn in SGC impaired axon regeneration, demonstrated by reduced axonal length and lower regeneration capacity (Fig. 5a-c). We also tested if loss of Fasn has an effect on the conditioning injury paradigm, in which a prior nerve injury increases the growth capacity of neurons ^82^. Isolated DRG neurons from naïve and injured FasncKO and control mice were cultured for 20h (Fig. 5d). In naïve animals, typically only few neurons extend short neurites, whereas a prior injury leads to more neurons initiating neurite growth and longer neurites (Fig. 5d-f). In FasncKO mice, naïve neurons presented similar neurite length to control naïve animals with a similar number of neurons initiating neurite growth. In contrast, a prior injury in FasncKO mice only partially conditioned the neurons for growth. Neurons in injured FasncKO displayed reduced neurite length compared to injured controls, but a similar number of neurons initiating neurite growth (Fig. 5d-f). These results indicate that Fasn in SGC contribute to the conditioning effect and the elongating phase of axon growth.

**Figure 5:**
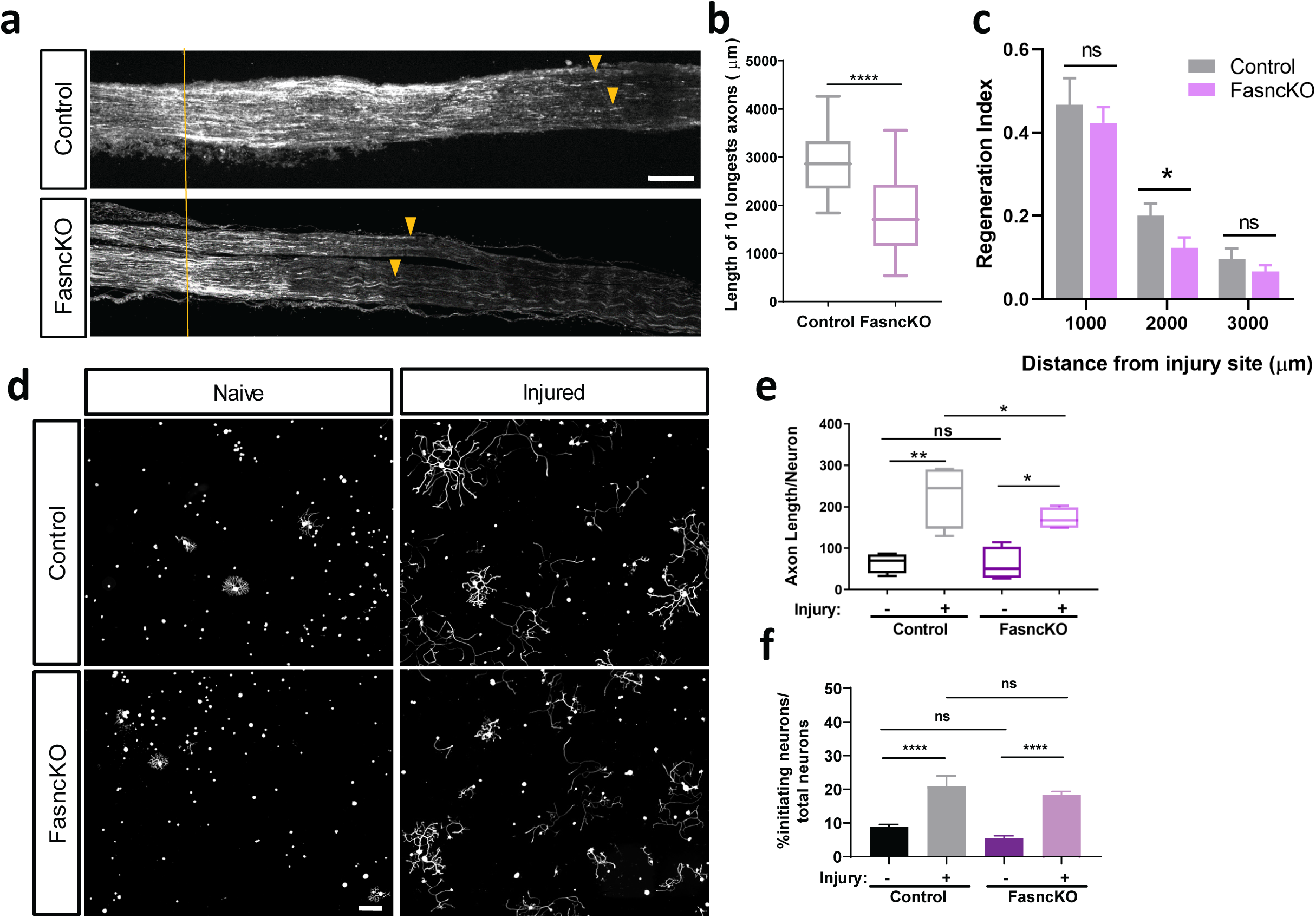
Fatty acid synthase deletion in SGC impairs axon regeneration. (a) Representative longitudinal sections of sciatic nerve from control and FasncKO mice 3 days after sciatic nerve injury, stained for SCG10. Orange lines indicate the crush site, identified as the maximal SCG10 intensity. Arrowheads indicate longest regenerating axons. Scale bar: 500µm (b) Length of the longest 10 axons was measured in 10 sections for each nerve. n=8. Unpaired t-test ****p<0.0001. Source data are provided as a Source Data file (c) Regeneration index was measured as SGC10 intensity normalized to the crush site. One way ANOVA. Sidak’s multiple comparisons test *p<0.05. Source data are provided as a Source Data file (d) Representative images of dissociated DRG neurons from control and FasncKO, cultured for 20h, from naïve and injured (3 days post conditioning sciatic nerve injury) and stained with the neuronal marker Tuj1. n=8. Scale bar: 200 µm. (e) Length of axons per neuron was measured. n=8, average of 500 neurons per replicate. Automated neurite tracing analysis using Nikon Elements. One way ANOVA. Sidak’s multiple comparisons test *p<0.05 **p<0.005 ns-non-significant. Source data are provided as a Source Data file (f) Percentage of initiating neurons normalized to the total number of neurons was measured n=8, average of 500 neurons per replicate. Automated neurite tracing analysis using Nikon Elements. One way ANOVA. Sidak’s multiple comparisons test ****p<0.0001 ns-non-significant. Source data are provided as a Source Data file.

### Activation of PPARα in SGC contributes to axon regeneration

To understand the mechanism by which Fasn in SGC regulates axon regeneration, we examined the role of peroxisome proliferator-activated receptors (PPARs), which represent a unique set of lipid regulated transcription factors ^34^. Fasn synthesizes palmitic acid, which is the substrate for the synthesis of more complex lipid species ^36^, including phospholipids. Importantly, it is not palmitic acid per se that is required for PPAR activation, but phospholipids ^37, 38^. Our scRNAseq analysis revealed that PPARα, but not PPARγ, was enriched in the SGC cluster, along with known PPARα target genes PPARGC1α, FADS2 and PEX11A ^83^ (Fig. 6a). These PPARα target genes were also upregulated after injury in the SGC cluster (Fig. 6b and Table 3,4). PPARα regulates the expression of genes involved in lipid and carbohydrate metabolism ^34^. This is consistent with our GO analysis of upregulated genes after injury that indicated an enrichment of the PPAR signaling pathway and lipid metabolism in SGC after injury (Fig. 3a and Table 4). We further confirmed the enrichment of PPARα target genes in SGC vs. neurons using an RNAseq data set from purified nociceptors in naïve condition and 3 days post injury that we previously generated ^84^ (Fig. 6b). These observations indicate that PPARα signaling is enriched in the SGC cluster and suggest that fatty acid synthesis in injured SGC regulates PPAR-dependent transcription.

**Figure 6:**
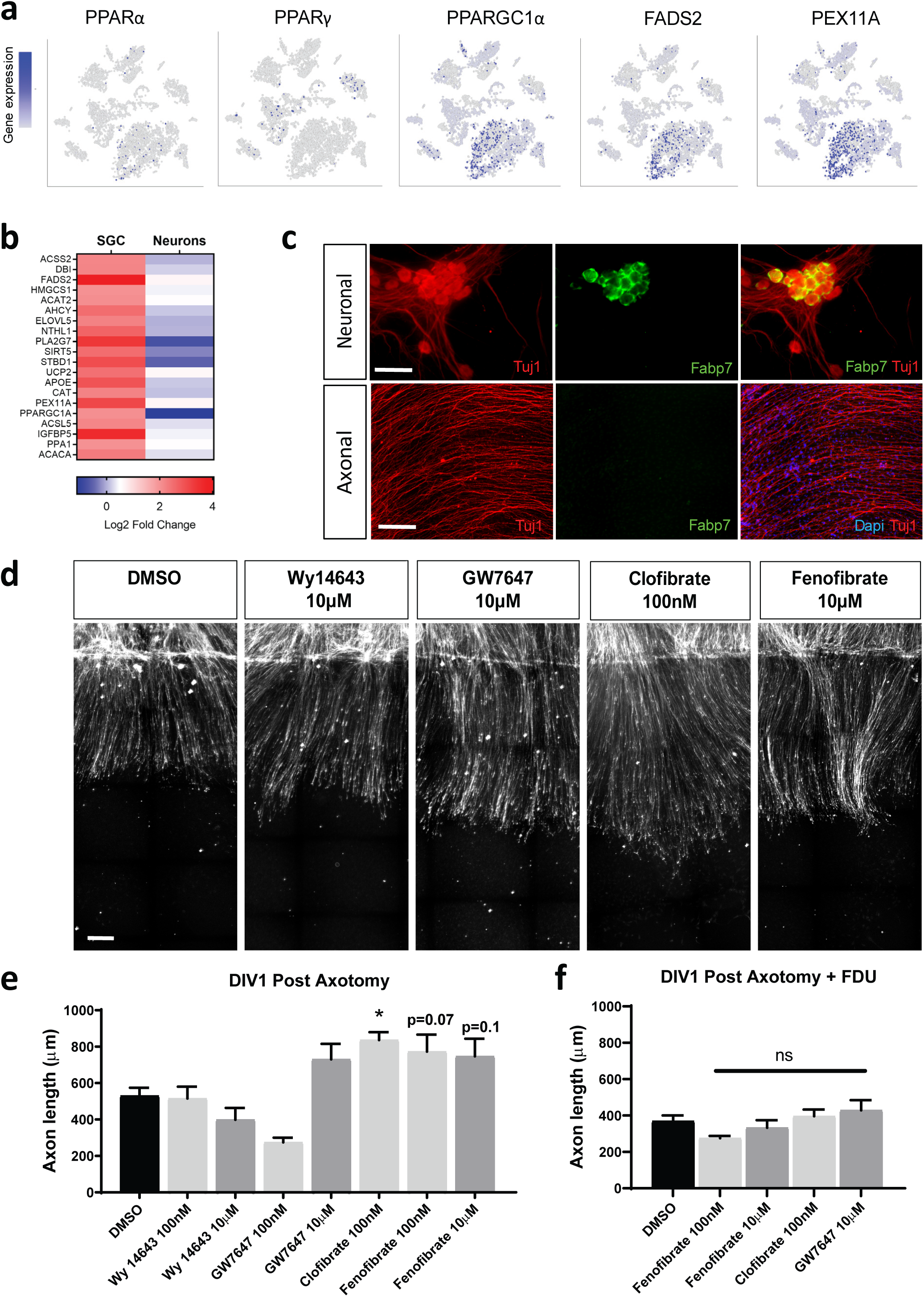
Activation of PPARα in SGC promotes axon regeneration in co-cultures of neurons and SGC. (a) t-SNE overlay for expression of the genes PPARα, PPARγ, and PPARα target genes PPARGC1a, FADS2 and PEX11a. (b) Heat map of PPARα target genes response to injury in SGC and neurons. (c) Neuronal and axonal area in embryonic DRG spot co-culture at DIV7, immunostained for Fabp7 (green) and Tuj1 (red). Scale Bar: 100 µm (neuronal) and 50 µm (axonal). (d) Embryonic DRG spot co-cultures axotomized at DIV7 after a 24 h pre-treatment with the indicated PPARα agonists at 2 concentrations (100nM and 10μM). Cultures were fixed after 24h and stained with SCG10. Scale Bar: 50 µm. (e) Distance of regenerating axons was measured from the injury site. n=4 One way ANOVA Dunnett’s multiple comparisons test. *p<0.05. Source data are provided as a Source Data file (f) Embryonic DRG spot neuronal culture was supplement with FDU (5-deoxyfluoruridine) to eliminate dividing cells in the culture. Axotomy was performed at DIV7 after a 24 h pre-treatment with the indicated PPARα agonists. Distance of regenerating axons was measured from injury site after 24h. n=4 One way ANOVA Dunnett’s multiple comparisons test. ns-non significant. Source data are provided as a Source Data file

Whereas biologic PPARα agonists consist of a broad spectrum of ligands ^83^, synthetic PPARα agonists include clofibrate, fenofibrate, bezafibrate, gemfibrozil, Wy14643 and GW7647. To test the effect of these PPARα agonists on axon regeneration, we modified our spot culture assay, in which embryonic DRG are dissociated and cultured in a spot, allowing axons to extend radially from the spot ^78, 80^. This assay recapitulates for the most part what can be observed *in vivo* in the nerve ^78, 79, 80^, and is thus suitable to test compounds affecting axon regeneration. By not including the mitotic inhibitor 5-deoxyfluoruridine (FDU) to eliminate dividing cells, we observed SGC, labeled with Fabp7 in the cell soma area, where they surround neurons by DIV7 (Fig. 6c, S5a,b) but not in the axon region (Fig. 6c, S5a,b). The PPARα agonists clofibrate, fenofibrate, Wy14643 and GW7647 were added to the media at 2 concentrations (10μM and 100nM) at DIV6. Axons were cut using a microtome blade at DIV7 and allowed to regenerate for 24h, fixed and stained for SCG10 to visualize axon regeneration ^81^. Regenerative length was measured from the visible blade mark to the end of the regenerating axon tips (Fig. 6d,e). Whereas Wy14643 and GW7647 had no effect on axonal regeneration compared to DMSO, clofibrate (100nM) and fenofibrate (100nM, 10μM) increased axon growth after axotomy (Fig. 6e). However, clofibrate at a higher concentration (10μM) caused cell death and axonal degeneration. To ensure that the increased regeneration effect was mediated by SGC and was not due to PPAR activation in neurons, we tested fenofibrate, clofibrate and GW7647 in spot cultures treated with FDU, in which no Fabp7 positive cells was detected at DIV7 (Fig. S5b). In these conditions, we observed no increase in axon regeneration (Fig. 6f and S5c). These results indicate that activation of PPARα signaling in SGC promote axon growth, although we cannot fully exclude that other cell types present in the spot culture contribute to this effect.

### Activation of PPARα with fenofibrate rescues the impaired axon regeneration in FasncKO mice

In the absence of Fasn in SGC, PPARα may lack its endogenous agonist and PPARα signaling following nerve injury in SGC may be compromised. To test this hypothesis, we determined if fenofibrate can rescue the axon regeneration defects that we observed in the FasncKO mice. We chose fenofibrate for these *in vivo* experiments because it is FDA-approved to treat dyslipidemia ^85^, is a specific PPARα agonist with minimal activity towards PPARγ and can be delivered easily in the diet ^86^. Control and FasncKO mice were fed with chow supplemented with fenofibrate, as described ^86^ for 2 weeks prior to nerve injury. PPARα as well as PPARα target genes were upregulated in the DRG of fenofibrate fed mice compared to mice fed with a normal diet (Fig. 7a). The reduced ATF3 expression in the DRG after nerve injury, in the FasncKO compared to control was rescued in the FasncKO mice fed with fenofibrate (Fig. 7b). Since ATF3 upregulation in response to injury occurs mainly in neurons (Fig. S6a), this results suggest that fenofibrate can compensate for the lack of Fasn in SGC and can rescue the neuronal response to injury. Another pro-regenerative neuronal gene, GAP43, showed a similar trend as ATF3. GAP43 reduced expression in response to injury in the FasncKO was rescued with the fenofibrate diet (Fig. S6b). In the FasncKO, we also noted an increase in GAP43 in naïve conditions with the fenofibrate diet compared to normal diet (Fig. S6b). This is consistent with previous reports that lipoproteins secreted from glial cells upregulate mRNA expression of GAP43 in hippocampal neurons ^42^. In contrast, JUN, which is also considered to be a pro-regenerative gene, was not affected in the FasncKO or by fenofibrate treatment (Fig. S6c). Because we assessed ATF3, Gap43 and Jun expression in whole DRG, we cannot exclude the possibility that other cell types regulate the expression of these genes in response to injury and fenofibrate treatment. We found that the impaired axon regeneration in FasncKO mice was rescued by the fenofibrate diet (Fig. 7c-e). The fenofibrate diet also rescued the impaired axon growth in the conditioning paradigm (Fig. 7f,g), with no change in the percent of neurons initiating growth (Fig. 7h). The regeneration of peripheral sensory neurons after injury is a relatively efficient prosses and we did not observe further improvement in both *in vivo* and *in vitro* experiment in control animals treated with fenofibrate (Fig. 7c-h). These results suggest that by activating PPARα, fenofibrate rescues the absence of Fasn in SGC. We cannot exclude the possibility that fenofibrate may also operate via other mechanisms on other cell types. However, these results are consistent with the notion that in response to axon injury, Fasn expression in SGC activates PPARα signaling to promote axon regeneration in adult peripheral nerves.

**Figure 7:**
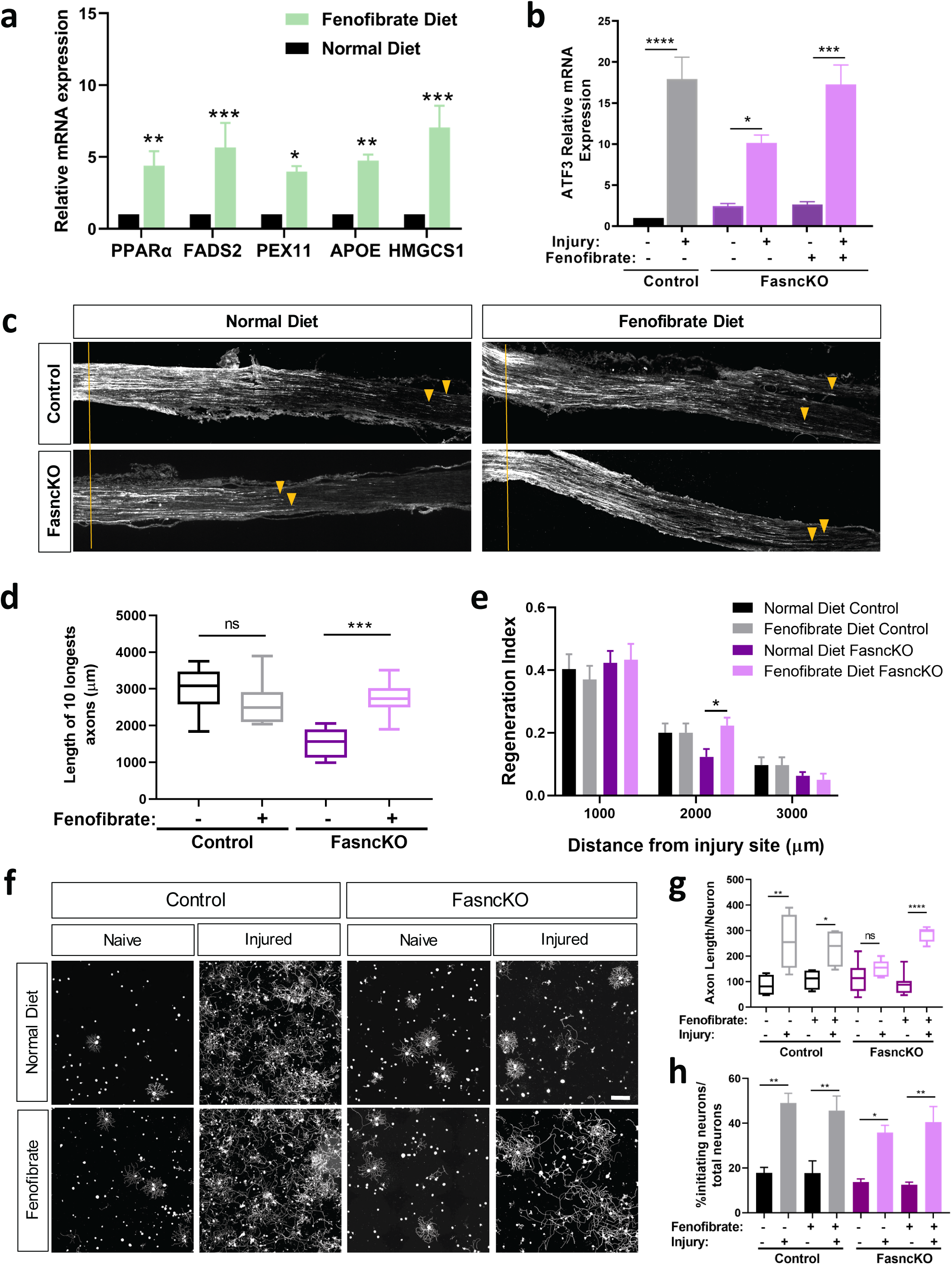
Activation of PPARα in SGC promotes axon regeneration. (a) qPCR analysis of PPARα target genes expression in DRG from mice fed with a normal diet vs. fenofibrate diet. n=3 One way ANOVA. Sidak’s multiple comparisons test *p<0.05 **p<0.005 ***p<0.0005. Source data are provided as a Source Data file (b) qPCR analysis of ATF3 expression in DRG from control and FasncKO mice in naïve and 3 days after sciatic nerve injury with and without fenofibrate n=3 One way ANOVA. Sidak’s multiple comparisons test *p<0.05 ***p<0.0005 ****p<0.0001. Source data are provided as a Source Data file (c) Representative longitudinal sections of sciatic nerve from FasncKO mice fed with a normal diet or fenofibrate diet, stained for SCG10. Orange lines indicate the crush site, arrow heads indicate longest axons Scale Bar: 500 µm. (d) Length of the longest 10 axons was measured in 10 sections for each nerve. n=8. One way ANOVA Sidak’s multiple comparisons test ***p<0.0005 ns-non significant. Source data are provided as a Source Data file (e) Regeneration index was measured as SGC10 intensity normalized to the crush site. One way ANOVA. Sidak’s multiple comparisons test *p<0.05. Source data are provided as a Source Data file (f) Representative images of dissociated DRG neurons from control and FasncKO mice fed with a normal diet or fenofibrate diet, cultured for 20h and stained with Tuj1. n=8. Scale bar: 200 µm. (g) Length of axons per neuron was measured. n=8, average of 500 neurons per replicate. One way ANOVA. Sidak’s multiple comparisons test *p<0.05 **p<0.005 ****p<0.0001 ns-non significant. Source data are provided as a Source Data file (h) Percentage of initiating neurons out of total number of neurons was measured n=8, average of 500 neurons per replicate. Automated neurite tracing analysis using Nikon Elements. One way ANOVA. Sidak’s multiple comparisons test *p<0.05 **p<0.005. Source data are provided as a Source Data file

## DISCUSSION

A role for SGC in nerve regeneration has not been considered and the biology of SGC is very poorly characterized under normal or pathological conditions. Our unbiased single cell approach characterized the molecular profile of SGC and demonstrated that SGC play a previously unrecognized role in peripheral nerve regeneration, in part via injury induced activation of PPARα signaling. Our results highlights that the neuron and its surrounding glial coat form a functional unit that orchestrates nerve repair.

SGC have been identified mostly based on their morphology and location. Several SGC markers have been characterized, but no useful markers that can be used to purify SGC and understand their biology at the molecular level have been identify so far. Although a previous study reported that SGC are very similar to Schwann cells ^11^, this conclusion was drawn from the analysis of cultured and passaged SGC and Schwann cells. Other SGC isolation methods rely on immunoreactivity to SK3 ^52^ or on multiple sucrose gradients and cell plating ^11^. One recent paper used immunisolation based on GS expression in SGC to examine their transicptonal profile and response to injury ^51^. Whereas at the immunostaining level, GS appear specifc to SGC, it does not appear to label all SGC ^51^. At the gene expression level, our data reveal that GS is expressed by many cell types in the DRG. Therefore immunoisolation based on GS expression might represent a subset of SGC and might not entirely be speciifc to SGC ^51^. Our single cell approach is unbaised to any predetermined markers and provides the advantage of minimal time between dissociation and analysis to capture more accurately the transcription status of the cells. Using this method, we demonstrate that SGC represent a unique cell population in the DRG, which share some molecular markers with Schwann cells as well as astrocytes.

We also characterized a new marker for adult SGC and a mouse line, BLBPcre-ER ^64^, that can be used to manipulate SGC specifically in the DRG. Because Fabp7 is expressed in radial glial cells as well as neuronal progenitors and is critical for neurogenesis in the CNS ^56, 87, 88^, this mouse line needs to be used in an inducible manner in adult mice to avoid targeting other cells during development. It is also important to note that Fabp7 is expressed in astrocytes, where it is important for dendritic growth and neuronal synapse formation ^89^ as well as for astrocyte proliferation during reactive gliosis ^90^, and thus the BLBPcre-ER line will also target astrocytes in the central nervous system. SGC are believed to share common features with astrocytes, such as expression of Kir4.1 and functional coupling by gap junctions ^7, 13, 16, 17^, with SGC surrounding the same neuron connected by gap junctions. The BLBPcre-ER mouse line thus represents a new tool to study SGC in the peripheral nervous system.

Injury related changes in SGC has been studied in pathological conditions such as inflammation, chemotherapy-induced neuropathic pain and nerve injuries ^7, 22, 23, 24^. Nerve injury was shown to increases SGC communication via gap junctions ^23^, leading to increased neuronal excitability ^24^. Nerve lesions also induce an increase in GFAP expression ^16, 22, 26, 27, 28, 29^. In our single cell data, we also found increased expression of connexin43/GJA1 and GFAP after nerve injury. Wheras prior studies also suggested SGC proliferation after injury ^7, 13, 16, 32^, a recent study demonstrated that cell proliferation occurred in macrophages but not in SGC after nerve injury ^51^. Our results are in agreement with this latter study, as we did observe enrichment for cell cycle markers in macrophages, and these cell cycle markers were downregulated in SGC 3 days after nerve injury. Prior studies relied largely on morphological location of cell cycle markers relative to neurons ^13, 16, 21^, and it is thus possible that macrophages infiltrating between neuron and SGC ^91^ were misidentified as proliferating SGC.

Our analysis revealed that SGC activate PPARα signaling to promote axon regeneration in adult peripheral nerves. PPARs are ligand-activated nuclear receptors with the unique ability to bind lipid signaling molecules and transduce the appropriate signals derived from the metabolic environment to control gene expression ^34^. Accordingly, PPARs are key regulators of lipid and carbohydrate metabolism. PPARs can sense components of different lipoproteins and regulate lipid homeostasis based on the need of a specific tissue ^35^. Three different PPAR subtypes are known; PPARα, PPARβ/δ and PPARγ ^35^. PPARγ is important in neurons for axon regeneration ^40^. However PPARγ is not expressed in SGC according to our scRNAseq data. Rather, our results indicate that PPARα is enriched in SGC and required for efficient nerve regeneration, at least at early time points after nerve injury. Whether SGC contribute to sustain regenerative growth until the target is reinnervated and whether different neuronal subtypes are differentially affected by PPARα signaling in SGC remains to be tested. Fenofibrate is a specific PPARα agonist, making it unlikely that the rescue effects we observed in FasncKO mice are due to activation of PPARγ in neurons. PPARα and PPARα target genes are also highly enriched in SGC, suggesting that fenofibrate acts on SGC rather than other resident cells in DRG to enhance axon regeneration. We also show that fenofibrate does not improve axon regeneration in neuronal cultures that do not contain SGC. However, we cannot exclude the possibility that fenofibrate may operate via other mechanisms in other cell types *in vivo*. Fenofibrate is used clinically to treat lipid disorders, but has unexpectedly shown in clinical trials to have neuroprotective effects in diabetic retinopathy ^92, 93^ and in traumatic brain injury ^94^. Our findings suggest that fenofibrate may be used to activate SGC and enhance axon regeneration in circumstances with poor sensory axon growth, such as after injury to dorsally projecting sensory axons in the spinal cord or peripheral nerve repair. Although fenofibrate treatment did not improve locomotor recovery following spinal contusion injury in mice ^95^, a slight trend for increased tissue sparing was observed. Whether fenofibrate can improve centrally projecting sensory axon growth will need to be rigorously tested.

Biologic PPARα agonists consist of saturated and unsaturated phospholipids, eicosanoids and glucocorticoids ^35^. Fasn synthesizes palmitic acid, which is the substrate for the synthesis of more complex lipid species ^36^. Fasn is partially localized in peroxisomes, where ether lipids such as plasmalogens are synthesized ^36^. In the liver, Fasn is required for generating the endogenous ligand for PPARα ^37^, whereas in adipose tissue, Fasn is required for generating endogenous ligands for PPARγ ^38^. Importantly, it is not palmitic acid per se that is required for PPAR activation but phospholipids. Phospholipid synthesis through the Kennedy pathway preferentially requires endogenous palmitate synthesis by Fasn rather than circulating palmitate from the diet ^37 38^. Our findings thus suggest that in SGC, Fasn generates ligands for PPARα following nerve injury and that Fasn dependent PPARα activation contribute to promote axon regeneration. Future experiments to conditionally delete PPARα from SGC may further strengthen the case that Fasn dependent PPARα activation in SGC after injury represent an important part of the injury response to stimulate axon repair.

How do lipid metabolism and PPARα signaling in SGC contribute to stimulate axon regeneration? Although addressing this question will require further investigation, our data indicate that Fasn expression in SGC participate in the regulation of a subset of regeneration associated gene after injury, including ATF3 and GAP43. This is consistent with previous reports that ApoE-containing lipoproteins secreted from glial cells leads to the upregulation of GAP43 mRNA in hippocampal neurons ^42, 48^. Together, these finding suggests that SGC contribute to regulate gene expression in neurons. It is also plausible that SGC impact neurons through lipid transfer. Indeed, lipids can be exogenously supplied by lipoproteins secreted from glial cells to stimulate neurite growth. Several studies have suggested a role for ApoE, a PPARα target gene, in lipid delivery for growth and regeneration of axons after nerve injury ^45, 47, 48^ and ApoE expression is increased in glial cells after nerve injury ^49, 50^. In agreement with these previous reports, our data indicate that ApoE mRNA levels are elevated in SGC after nerve injury and also following fenofibrate treatment. A recent study described downregulation of few genes related to cholesterol metabolism after injury in SGC, but whether and how this contribute to axon repair has not been evaluated ^51^. Cholesterol depletion was also shown to improve axon regeneration ^96^, which contrasts with earlier studies showing that cholesterol is required for axonal growth and can be synthesized in neurons or exogenously supplied from lipoproteins to axons of cultured neurons.^47^. It will therefore be interesting to determine if lipid metabolism affecting cholesterol, fatty acids and phospholipids in SGC regulates axon regeneration through paracrine effects on neurons.

Our study suggests that what has been defined as a neuronal intrinsic response to injury, namely upregulation of regeneration associated genes, such as ATF3 and GAP43 in neuron, ^1, 2, 97^ may depend in part on signaling from SGC surrounding the neuronal soma. The neuron and its surrounding glial coat may thus form a functional unit that orchestrates nerve repair. Our single cell data set also highlights that other cell types, including endothelial cells, mesenchymal cells and macrophages in the DRG respond to a distant nerve injury. Future studies are needed to elucidate the complex cellular cross talks operating in the DRG after nerve injury that contribute to nerve repair.

## ACKNOWLEDGMENTS

We would like to thank members of the Cavalli lab for valuable discussions. We would like to thank the following investigators for their generous gifts of mouse lines: Dr. Toshihiko Hosoya for the BLBP-cre ER mice, Dr. Yuji Oawada for the Fabp7KO mice and Dr. Harrison Gabel for the Sun1GFP mice. We gratefully acknowledge Greg Strout, Ross Kossina and Dr. James Fitzpatrick from the Washington University Center for Cellular Imaging (WUCCI), which is supported in part by Washington University School of Medicine, The Children’s Discovery Institute of Washington University, and St. Louis Children’s Hospital (CDI-CORE-2015-505 and CDI-CORE-2019-813) and the Foundation for Barnes-Jewish Hospital (3770) for assistance in acquiring and interpreting Transmission Electron Microscopy (TEM) data. We also thank Anushree Seth and Madison Mack in association with InPrint for illustration in Fig.1a. This work was funded in part by a post-doctoral fellowship from The McDonnell Center for Cellular and Molecular Neurobiology to O.A, by NIH grant R35 NS111596 to V.A.K, and by The McDonnell Center for Cellular and Molecular Neurobiology and NIH grant NS111719 to V.C.

## AUTHOR CONTRIBUTIONS

O.A, P.D., R.K, C.F.S., V.A.K. and VC designed research as well as edited and approved the manuscript; O.A. performed sequencing, bioinformatics, *in vitro* and *in vivo* experiments, analyzed the data and wrote the manuscript; P.D. performed electrophysiological recordings experiments; S.J. analyzed and quantified in vitro and in vivo regeneration assays and TEM images.

## DECLARATION OF INTERESTS

The authors declare no competing interests.

## Supplementary Figures

**Figure S1:**
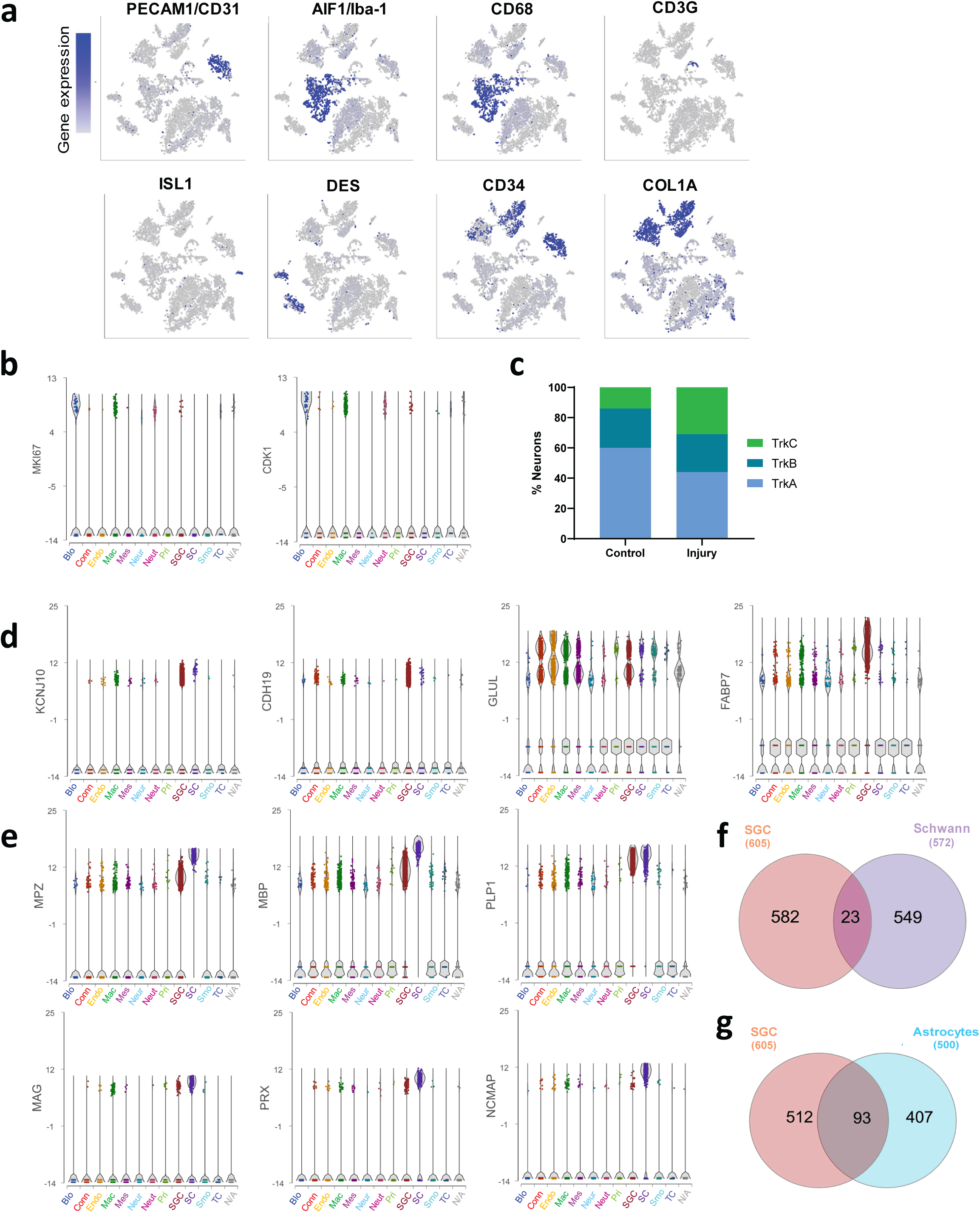
Cluster analysis from scRNAseq of mouse DRG, related to Figure 1. (a) t-SNE overlaid for expression of marker genes for different cell populations including PECAM/DC31 for endothelial cells, AIF1/Iba-1 and CD68 for Macrophages. CD3G for T-cells, Isl1 for neurons, DES for smooth muscle cells, CD34 for mesenchymal cell and COL1A1 for connective tissue. (b) Plots illustrate the gene counts (log) for Miki67 and Cdk1 genes of distinct cell populations in injury conditions. Color scheme for populations on the left. (c) Fraction of neuronal type within control and injury condition by expression of Trk receptors. Source data are provided as a Source Data file (d) Violin plots illustrate SGC marker genes signatures of distinct cell populations. (e) Violin plots illustrate Schwann and SGC marker genes signatures of distinct cell populations. (f) Top differentially expressed genes in the SGC cluster (605 genes) was compared to top differentially expressed genes in the Schwann cell cluster (572 genes) (g) Top differentially expressed genes in the SGC cluster (605 genes) was compared to the top differentially expressed genes in astrocytes.

**Figure S2:**
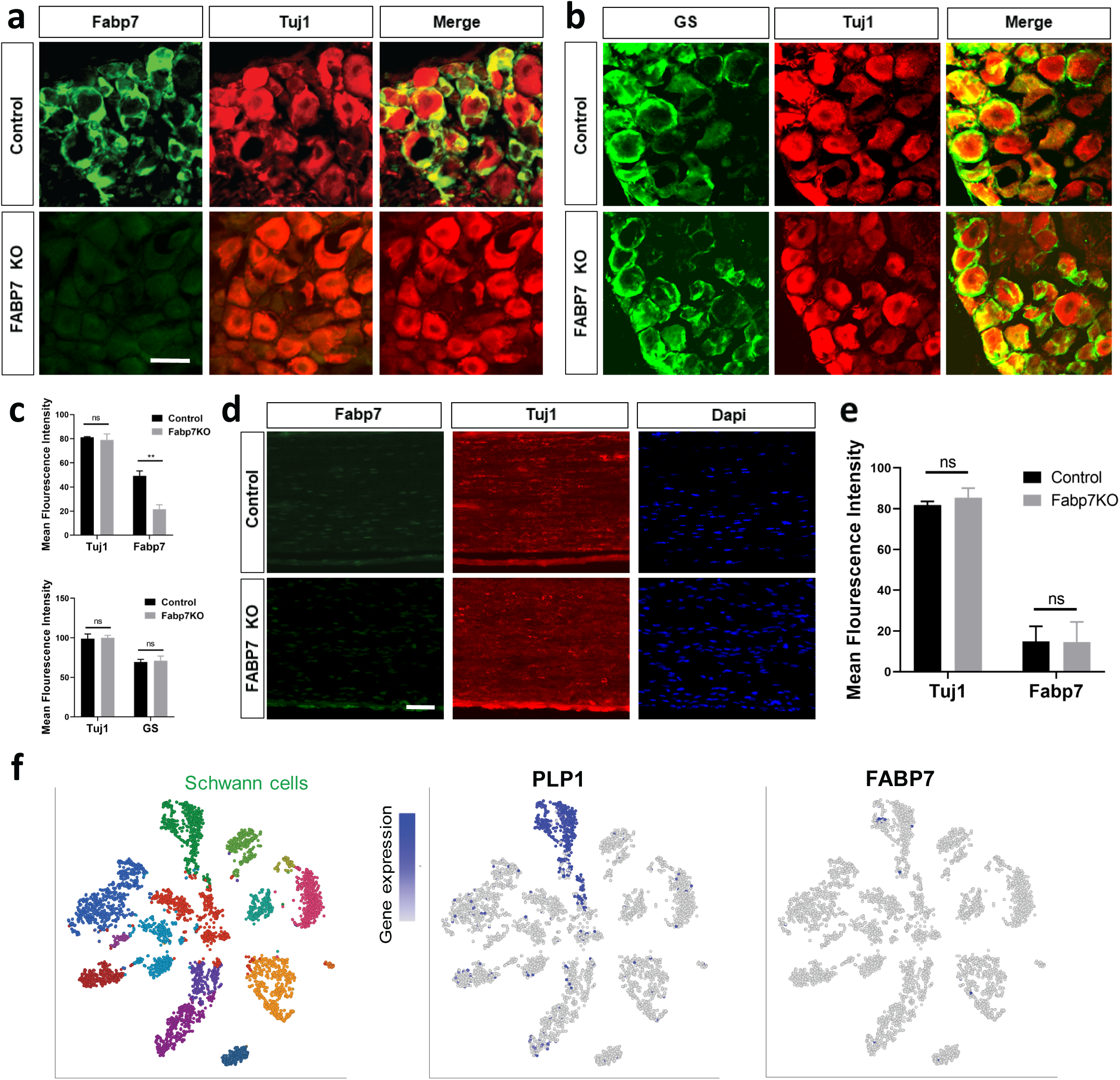
Fabp7 is a specific marker for SGCs in adult mice DRG, related to Figure 2. (a) DRG sections from Fabp7KO and control mice were immunostained for Fabp7 (green) and Tuj1 (red) (b) DRG sections from Fabp7KO and control mice were immunostained for Glutamine Synthase (GS) (green) and Tuj1 (red). Scale bar: 50 µm (c) Quantification of DRG mean fluorescence intensity of Fabp7/GS and Tuj1. Two-way ANOVA **p<0.005 ns-non significant. Source data are provided as a Source Data file (d) Longitudinal sections of sciatic nerves from Fabp7KO and control mice immunostained for Fabp7 (green), Tuj1 (red) and Dapi. Scale bar: 50 µm. (e) Quantification of nerves mean fluorescence intensity of Fabp7 and Tuj1. Two-way ANOVA ns-non significant. Source data are provided as a Source Data file (f) t-SNE plot of injured sciatic nerve (9 days post injury) analyzed from the single cell data (Carr et al., 2018). t-SNE overlay for the expression of PLP1 gene, indicating the Schwann cell cluster, and Fabp7.

**Figure S3:**
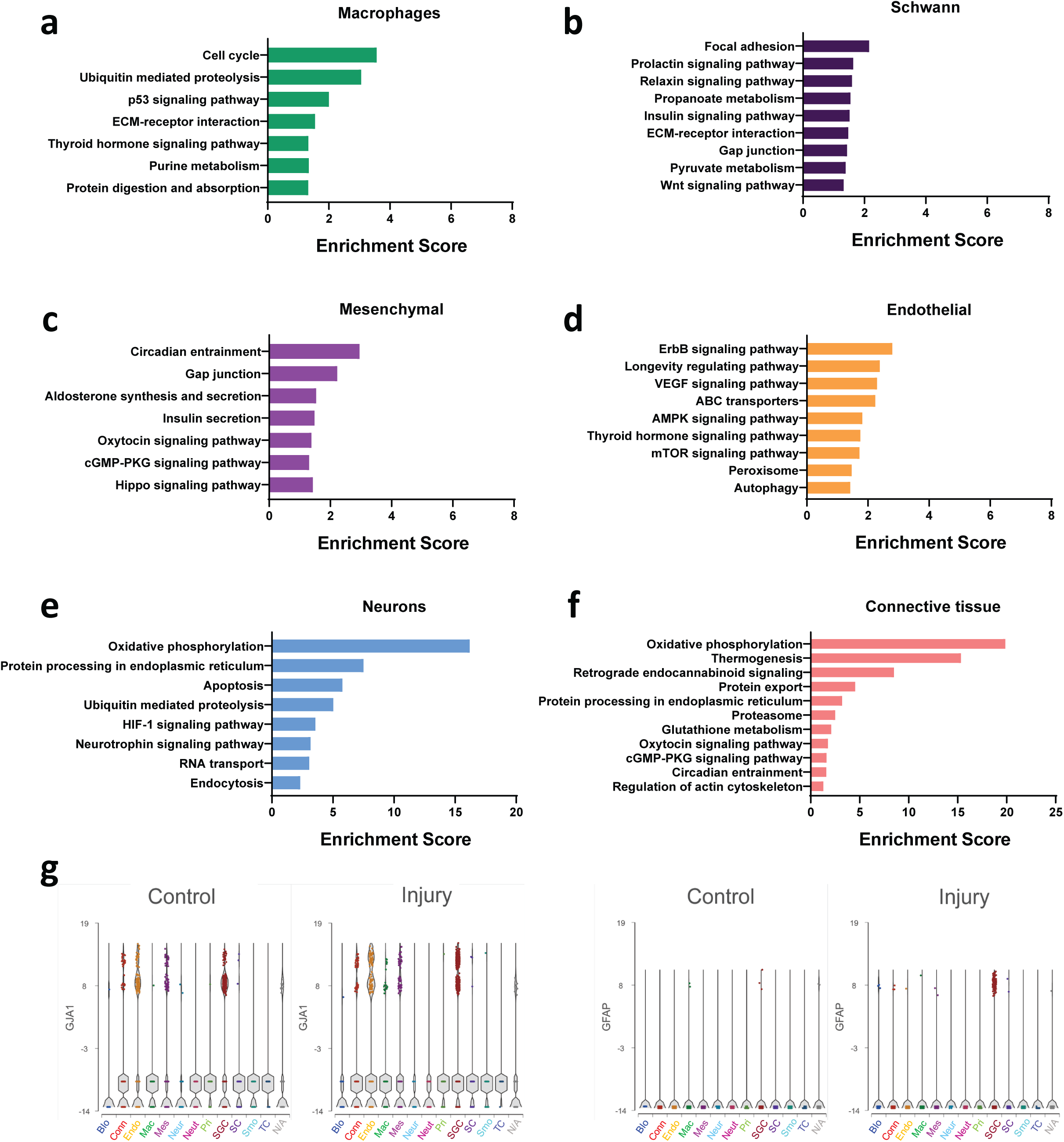
Pathway analysis of differentially upregulated genes in major cell types, related to Figure 3. (a-f) Pathway analysis of differentially upregulated genes in major cell types in the DRG following nerve injury (KEGG 2016). (g) Violin plots illustrate SGC injury induced genes GJA1 and GFAP signatures of distinct cell populations in control and injury conditions

**Figure S4:**
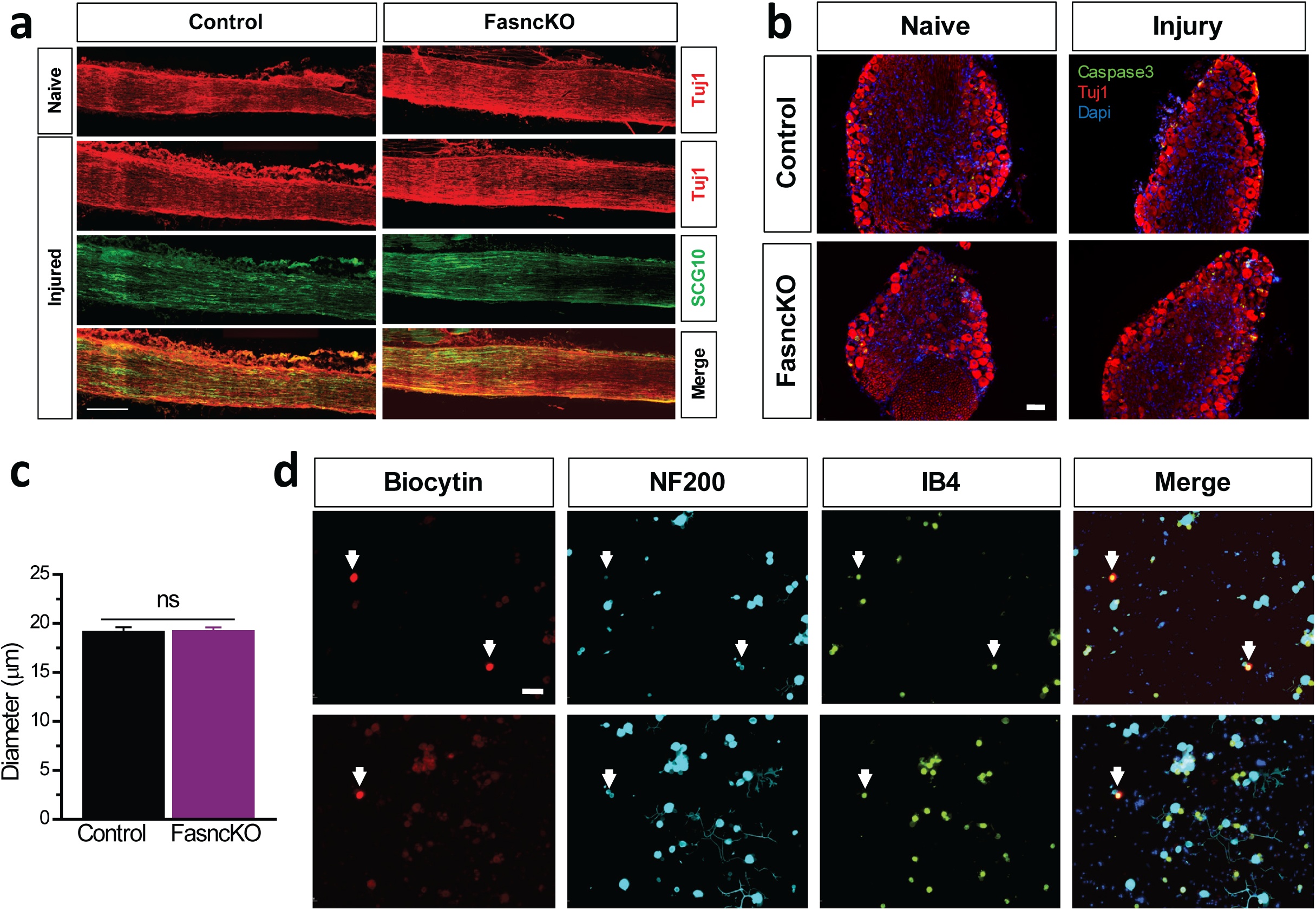
Fatty Acid synthase deletion in SGC does not lead to neuronal cell death or abnormal functional properties, related to Figure 4. (a) Representative images of longitudinal nerve sections from naïve and injured control and FasncKO mice, immunostained for TUJ1(red) and SCG10(green). Scale bar: 500 µm. (b) DRG sections from FasncKO and control mice in naïve and injured (3 days post injury) were immuonostained for Cleaved Caspase3 (green), Tuj1 (red) and Dapi (Blue). Scale bar: 50 µm. (c) Whole-cell recordings in dissociated co-cultures of DRG neurons and glia. Medium diameter neurons (control 19.19 ± 0.42 µm, n = 16; FasncKO 19.27 ± 0.34 µm, n = 30, p = 0.88; that were associated with at least one SGC, were targeted for recordings. Source data are provided as a Source Data file (d) A subset of recorded cells was filled with biocytin (red) via the patch pipette for post hoc verification of neuronal identity; IB4-nociceptors (green) NF200-Proprioceptors/LTMRs (Cyan). Scale bar: 50 µm.

**Figure S5:**
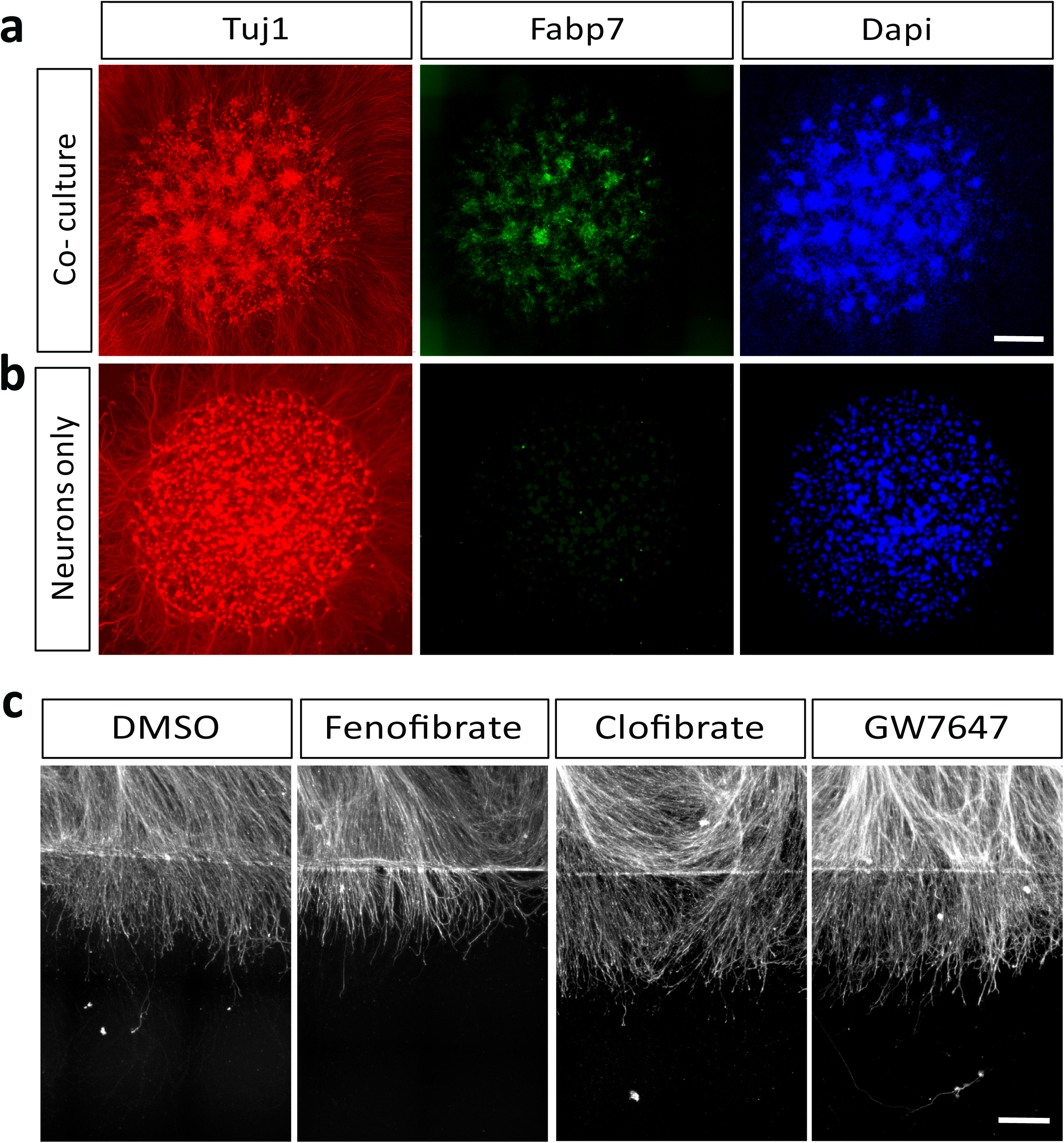
Activation of PPARα in pure neuronal cultures does not enhance axon regeneration, related to Figure 6. (a) Embryonic DRG were dissociated and plated as a spot without 5-fluorodeoxyuridine (FDU) and stained for Fabp7 (green), Tuj1 (red) and dapi (blue) at DIV6. n=5. (b) Embryonic DRG were dissociated and plated as a spot with 5-fluorodeoxyuridine (FDU) (c) Embryonic DRG spot co-culture, supplemented with FDU, were axotomized at DIV7 after a 24 h pre-treatment with the indicated PPARα agonists fenofibrate (10μM), Clofibrate (100nM) and GW7647 (10μM). Cultures were fixed after 24h and stained with SCG10. Scale Bar: 250 µm.

**Figure S6:**
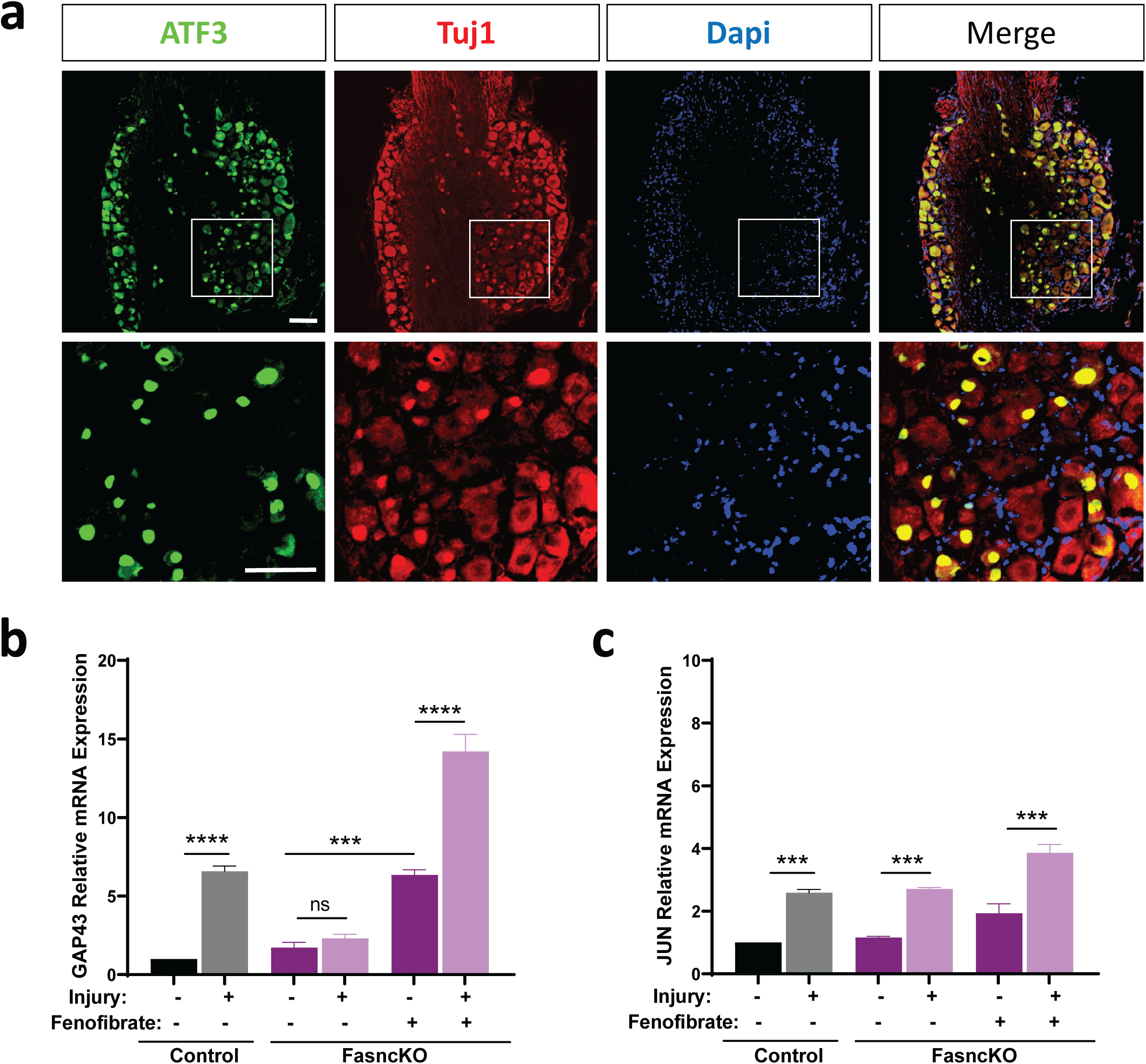
Neuronal pro-regeneration genes expression in response to fenofibrate, related to figure 7. (a) Representative images of injured DRG (3 days post injury) immunostained for ATF3(green), Tuj1(red) and Dapi(Blue) Scale Bar: 100 µm. (b) qPCR analysis of GAP43 expression in DRG from control and FasncKO mice in naïve and 3 days after sciatic nerve injury with and without fenofibrate n=3 One way ANOVA. Sidak’s multiple comparisons test ***p<0.001, ****p<0.0001 ns-non significant. Source data are provided as a Source Data file (c) qPCR analysis of JUN expression in DRG from control and FasncKO mice in naïve and 3 days after sciatic nerve injury with and without fenofibrate n=3 One way ANOVA. Sidak’s multiple comparisons test ***p<0.0005. Source data are provided as a Source Data file

## Supplementary Tables

**Table 1**: Top 10 expressed genes in each cell cluster

**Table 2:** Comparisons of genes expressed in SGC, Schwann cells, astrocytes and myelinating/non myelinating Schwann cells

**Table 3:** Differentially expressed genes in SGC after nerve injury

**Table 4**: Enriched up regulated pathways after injury in SGC with the corresponding genes.

## Material and Methods

### Animals and Procedures

All surgical procedures were completed as approved by Washington University in St. Louis School of Medicine Institutional Animal Care and Use Committee’s regulations. During surgery, 8-12 week old C57Bl/6 mice of the indicate genotype were anesthetized using 2% inhaled isoflurane. For analgesia, 1mg/kg buprenorphine SR-LAB (ZooPharm) was administered subcutaneously. Sciatic nerve injuries were performed as previously described ^80, 84^. Briefly, the sciatic nerve was exposed and crushed for 10 seconds using a #55 forceps. The wound was closed using wound clips and both injured L4 and L5 dorsal root ganglia and sciatic nerve were dissected at the indicated time post-surgery. Contralateral nerve and DRG served as uninjured controls, when needed. Tamoxifen (500mg per kg diet, TD.130858) and Fenofibrate (0.2%, Sigma Cat# F6020) were administrated as chow pellets (Envigo Teklad).

### Mouse strains

8-12 weeks old male and female mice were used for all experiments, except for scRNAseq experiment, where only C57Bl/6 females were used *Rosa26-ZsGreen* (also known as Ai6(RCL-ZsGreen) was obtained from Jackson RRID:IMSR_JAX:007906; ^98^. Fabp7KO mouse line was a generous gift from Dr. Owada ^61^. The Sunf1GFP mice (Gt(ROSA)26Sortm5(CAG-Sun1/sfGFP)Nat) was a generous gift from Dr. Harrison Gabel. mice carrying floxed *Fasn* alleles were previously generated ^70^. The BLBPcre-ER mouse line ^64^ was a generous gift from Dr. Toshihiko Hosoya.

### Single cell RNAseq

L4 and L5 DRG’s from mice 8-12 weeks old were collected into cold Hank’s balanced salt solution (HBSS) with 5% Hepes, then transferred to warm Papain solution and incubated for 20 min in 37°C. DRG’s were washed in HBSS and incubated with Collagenase for 20 min in 37°C. Ganglia were then mechanically dissociated to a single cell suspension by triturating in culture medium (Neurobasal medium), with Glutamax, PenStrep and B-27. Cells were washed in HBSS+Hepes +0.1%BSA solution, passed through a 70-micron cell strainer. Hoechst dye was added to distinguish live cells from debris and cells were FACS sorted using MoFlo HTS with Cyclone (Beckman Coulter, Indianapolis, IN). Sorted cells were washed in HBSS+Hepes+0.1%BSA solution and manually counted using hemocytometer. Solution was adjusted to a concentration of 500cell/microliter and loaded on the 10X Chromium system. Single-cell RNA-Seq libraries were prepared using GemCode Single-Cell 3′ Gel Bead and Library Kit (10x Genomics). A digital expression matrix was obtained using 10X’s CellRanger pipeline (Washington University Genome Technology Access Center). Quantification and statistical analysis were done with Partek Flow package (Build version 9.0.20.0417). Low quality cells and potential doublets were filtered out from analysis using the following parameters; total reads per cell: 600-15000, expressed genes per cell: 500-4000, mitochondrial reads <10%. A noise reduction was applied to remove low expressing genes <=1 count. Counts were normalized and presented in logarithmic scale in CPM (count per million) approach. An unbiased clustering (graph based clustering) was done and presented as t-SNE (t-distributed stochastic neighbor embedding) plot, using a dimensional reduction algorithm that shows groups of similar cells as clusters on a scatter plot. Differential gene expression analysis performed using an ANOVA model; a gene is considered differentially-expressed (DE) if it has an false discovery rate (FDR) step-up (p-value adjusted).*p* ≤ 0.05 and a Log2fold-change ≥±2. The data was subsequently analyzed for enrichment of GO terms and the KEGG pathways using Partek flow pathway analysis. Partek was also used to generate figures for t-SNE and scatter plot representing gene expression.

### Tissue Preparation and Immunohistochemistry

After isolation of either sciatic nerve or DRG, tissue was fixed using 4% paraformaldehyde for 1 hour at room temperature. Tissue was then washed in PBS and cryoprotected using 30% sucrose solution at 4C overnight. Next, the tissue was embedded in O.C.T., frozen, and mounted for cryosectioning. All frozen sections were cut to a width of 12µm for subsequent staining. For immunostaining of DRG and nerve sections, slides were washed 3x in PBS and then blocked for in solution containing 10% goat serum in .2% Triton-PBS for 1 hour. Next, sections were incubated overnight in blocking solution containing primary antibody. The next day, sections were washed 3x with PBS and then incubated in blocking solution containing a secondary antibody for 1 hour at room temperature. Finally, sections were washed 3x with PBS and mounted using ProLong Gold antifade (Thermo Fisher Scientific). Images were acquired at 10x or 20x using a Nikon TE2000E inverted microscope and images were analyzed using Nikon Elements. Antibodies were as follow: SCG10/Stmn2 (1:1000; Novus catalog #NBP1-49461, RRID:AB_10011569), Tubb3/βIII tubulin antibody (BioLegend catalog #802001, RRID:AB_291637), *Griffonia simplicifolia* isolectin B4 (IB4) directly conjugated to Alexa Fluor 488 or Alexa Fluor 594 (Thermo Fisher Scientific catalog #I21411 and #I21413), Fabp7 (Thermo Fisher Scientific Cat# PA5-24949, RRID:AB_2542449), cleaved caspase 3 (CST Cat# 9664, RRID:AB_2070042), Fasn (Abcam, Catalog #ab128870), Glutamine synthase (Abcam, Catalog #ab49873). Stained sections with only secondary antibody were used as controls.

### DRG Cultures and Regeneration Assays

For in vitro regeneration assay, dorsal root ganglia were isolated from time pregnant e13.5 CD-1 mice and cultured as previously described ^80, 99^. Briefly, after a short centrifugation, dissection media was aspirated and cells were digested in .05% Trypsin-EDTA for 25 minutes in 37°C. Next, cells were pelleted by centrifuging for 2 minutes at 500 x g, the supernatant was aspirated, and Neurobasal was added. Cells were then triturated 25x and added to the growth medium containing Neurobasal media, B27 Plus, 1ng/ml NGF, Glutamax and Pen/Strep, with or without 5μM 5-deoxyfluoruridine (FDU). Approximately 10,000 cells were added to each well in a 2.5 μl spot. Spotted cells were allowed to adhere for 10 minutes before the addition of the growth medium. Plates were pre-coated with 100ug/ml poly-D-lysine. For regeneration assays, PPARα agonists were added to the culture on DIV6. Cells were then injured using an 8mm microtome blade on DIV7 and fixed 24h later. Cells were washed with PBS and stained for SCG10 as described above.

For adult DRG cultures DRG were dissected from naïve mice. Cells were prepared as described above for single cell protocol and cultured on 100ug/ml poly-D-lysine coated plates and fixed 20h later. Cultures were then used for electrophysiological recording 24h after plating or fixed and stained with the indicated antibody. Images were acquired at 10x using a Nikon TE2000 microscope and image analysis was completed using Nikon NIS-Elements (Version 4.60).

### Image Analysis

For sciatic nerve injury experiments, images were quantified in two ways. First, the injury site was defined as the area with maximal SCG10 intensity, as described previously ^79, 100^. A vertical line was drawn across this region and the longest 10 axons were measured from that site and the average length reported, as described ^84^. Next, SCG10 intensity was quantified at 1, 2, and 3mm from the injury site as previously described ^79, 80^. The intensity was normalized to the injury site and the percent intensity was reported. For both length and intensity quantifications, five sections per biological replicate were averaged.

For embryonic dorsal root ganglia experiments, regenerative length was measured from the visible blade mark to the end of the regenerating axons. Each technical replicate was measured 4-6 times and three technical replicates were measured per biological replicate.

To determine the cleaved caspase staining area, a binary was generated to fit the positive signal, and positive staining area was measured. That area was internally normalized to Dapi positive staining area.

### RNA Isolation and Quantitative PCR

DRG and nerves were lysed and total RNA was extracted using Trizol reagent (Thermo Fisher, Cat# 15596026).). Next, RNA concentration was determined using a NanoDrop 2000 (Thermo Fisher Scientific). First strand synthesis was then performed using the High Capacity cDNA Reverse Transcription kit (Applied Biosystems). Quantitative PCR was performed using PowerUp SYBR Green master mix (Thermo Fisher, Cat# a25742) using 5ng of cDNA per reaction. Plates were run on a QuantStudio 6 Flex and analyzed in Microsoft Excel. The average Ct value from three technical replicates was averaged normalized to the internal control Rpl13a. All primer sequences were obtained from PrimerBank ^101^ and product size validated using agarose gel electrophoresis.

Pparα (PrimerBank ID 31543500a1) Forward Primer AGAGCCCCATCTGTCCTCTC Reverse Primer ACTGGTAGTCTGCAAAACCAAA

Pex11a (PrimerBank ID 6755034a1) Forward Primer GACGCCTTCATCCGAGTCG Reverse Primer CGGCCTCTTTGTCAGCTTTAGA.

Fads2 (PrimerBank ID 9790070c1) Forward Primer TCATCGGACACTATTCGGGAG Reverse Primer GGGCCAGCTCACCAATCAG.

Hmgcs1 (PrimerBank ID 31981842a1) Forward Primer AACTGGTGCAGAAATCTCTAGC Reverse Primer GGTTGAATAGCTCAGAACTAGCC

ApoE (PrimerBank ID 6753102a1) Forward Primer CTGACAGGATGCCTAGCCG Reverse Primer CGCAGGTAATCCCAGAAGC

ATF3 (PrimerBank ID 31542154a1) Forward Primer GAGGATTTTGCTAACCTGACACC Reverse Primer TTGACGGTAACTGACTCCAGC

Gap43 (PrimerBank ID 6679935a1) Forward Primer TGGTGTCAAGCCGGAAGATAA Reverse Primer GCTGGTGCATCACCCTTCT

Jun (PrimerBank ID 6680512a1) Forward Primer TCACGACGACTCTTACGCAG Reverse Primer CCTTGAGACCCCGATAGGGA

Rpl13a (PrimerBank ID 334688867c2) Forward Primer AGCCTACCAGAAAGTTTGCTTAC Reverse Primer GCTTCTTCTTCCGATAGTGCATC

### Electrophoresis and western blot

DRG and sciatic nerve samples were lysed in 5× SDS loading buffer, heated at 95 °C for 5 min and were run in on 8–16% NuPAGE SDS-PAGE gradient gel (Life Technologies) in MOPS SDS Running Buffer (Life Technologies) and transferred to 0.45 μm nitrocellulose membranes (Amersham). Next, these membranes were blocked in blocking buffer (3% BSA in 0.1% Tween-20 at pH 7.6 for 1 h, and incubated at 4 °C overnight in blocking buffer with primary antibodies. Membranes were then washed in TBST 3 times, incubated with horseradish peroxidase (HRP)-conjugated anti-rabbit or anti-mouse IgG (Invitrogen) for 1 h at room temperature, washed in TBST three times, and developed with SuperSignal West Dura Chemiluminescent Substrate (Thermo Scientific). The membranes were imaged using ChemiDoc System (Bio Rad) and analyzed with ImageLab software. Gapdh (Santa Cruz, catalog# sc25778) was used as loading control for quantification of the protein blots, Fasn intensity was normalized to the intensity of Gapdh.

### Whole-cell electrophysiology

Whole-cell patch-clamp recordings in a current-clamp mode were performed using a Multiclamp 700B amplifier (Molecular Devices) from short-term cultures (24 hours after plating)isolated DRG neurons visually identified with infrared video microscopy and differential interference contrast optics (Olympus BX51WI). Current-clamp recordings were made with pipette capacitance compensation and bridge-balance compensation. Recordings were conducted at near-physiological temperature (33–34°C). The recording electrodes were filled with the following (in mM): 130 K-gluconate, 10 KCl, 0.1 EGTA, 2 MgCl2, 2 ATPNa2, 0.4 GTPNa, and 10 HEPES, pH 7.3. The extracellular solution contained (in mM): 145 NaCl, 3 KCl, 10 HEPES, 2.5 CaCl2, 1.2 MgCl2, and 7 glucose, pH 7.4 (saturated with 95% O2 and 5% CO2). For determination of action potential (AP) threshold, APs were evoked by a ramp current injection (0.1 pA/ms) ^102, 103^ with a hyperpolarizing onset to ensure maximal Na^+^ channel availability before the first AP. The AP thresholds were determined only from the first APs of ramp-evoked AP trains. AP threshold (i.e., threshold voltage) was defined as the voltage at the voltage trace turning point, corresponding to the first peak of 3rd order derivative of AP trace ^102, 104^. Data were averaged over 5-8 trials for each cell. Resting membrane potential (RMP) was measured immediately after whole-cell formation. Cell capacitance was determined by the amplifier’s auto whole-cell compensation function with slight manual adjustment to optimize the measurement if needed. Under current-clamp mode, a negative current (−50 pA for 500 ms) was injected every 5 s to assess the input resistance.

### TEM

Mice were perfused with 2.5% glutaraldehyde with 4% paraformaldehyde in 0.1M Cacodylate buffer, followed by post fix. A secondary fix was done with 1% osmium tetroxide. For Transmission electron microscopy (TEM), tissue was dehydrated with ethanol and embedded with spurr’s resin. Thin sections (70 nm) were mounted on mesh grids and stained with 8% uranyl acetate followed by Sato’s lead stain. Sections were imaged on a Jeol (JEM-1400) electron microscope and acquired with an AMT V601 digital camera. (Washington University Center for Cellular Imaging).

### Quantification and statistical analysis

Quantifications were performed by a blinded experimenter to genotype and treatment. Fiji (ImageJ) analysis software was used to measure TEM nerve images. NIkon Elements analysis software was used to trace regenerating axon in nerve sections and in the embryonic DRG spot culture. An Automated analysis for axon tracing and neurons soma count was used for ex-vivo adult DRG culture experiments (Nikon elements commercial software package, code is available upon request). Statistics was performed using GraphPad (Prism8) for either Student’s t-test or ANOVA analysis. Sidak or Dunnett tests were used as part of one- and two-way ANOVA for multiple comparison tests. Error bars indicate the standard error of the mean (SEM).

### Data Availability

The Fastq files and the filtered count matrix for scRNA sequencing were deposited at the NCBI GEO database under the accession number GSE139103.

Data analysis and processing was performed using commercial code from Partek Flow package https://www.partek.com/partek-flow/

Axon tracing and neurons soma count was performed using Nikon NIS-elements, which is a commercial software package for image analysis. The specific analysis code for digital reconstruction of axons is available upon request (requires NIS-Elements and General Analysis) https://www.microscope.healthcare.nikon.com/products/software/nis-elements

The source data file contain the raw data underlying all reported averages in graphs and charts, and uncropped version of blot presented in figures 1e,2c,3e-f,f,4b-d,f-q,5b,c,e,f, 6e-f, 7a-b,d-e,g-f and in supplemental figures S1c,S2c,e, S4c, S6b-c.

## Notes

### Competing Interest Statement

The authors have declared no competing interest.

